# Regulating the timing of enhancer transitions is key to defining sharp boundaries of Fushi tarazu expression in the *Drosophila* embryo

**DOI:** 10.1101/2022.09.27.509025

**Authors:** Anthony Birnie, Audrey Plat, Jacques P. Bothma

## Abstract

Coordinating the action of different enhancers is crucial to correctly specify cell fate decisions during development. Yet it remains poorly understood how the activity of multiple enhancers is choregraphed in time. To shed light on this question we used new live imaging approaches to quantify transcription and protein expression in single cells of *Drosophila melanogaster* embryos. We employed these tools to dissect the regulation of Fushi tarazu (Ftz), a transcription factor expressed in a series of stripes by two distinct enhancers: autoregulatory and zebra. The anterior edges of the Ftz stripes are sharply defined and specify essential signaling centers. Here, we determined the time at which each boundary cell commits to either a high-Ftz or low-Ftz fate using dynamic features of time-resolved Ftz protein traces. By following the activity of each enhancer individually, we showed that the autoregulatory enhancer does not establish this fate choice. Instead, it perpetuates the decision defined by zebra. This is contrary to the prevailing view that autoregulation drives the fate decision by causing bi-stable Ftz expression. Furthermore, we showed that the autoregulatory enhancer is not activated based on a Ftz concentration threshold, but through a timing-based mechanism. We hypothesize that this is regulated by a set of pioneer-like transcription factors, which have recently been shown to act as timers in the embryo. Our work provides new insight into the genetic mechanisms that directly regulate the dynamics of gene regulatory networks, and supports the emerging view that this regulation is vital for reliable cell fate specification.

## Introduction

The genes that direct cellular differentiation often need to change which biological pathways they respond to^1^. This is governed by the multiple regulatory regions of DNA, called enhancers, that typically control the expression of an individual gene^2,3^. Different enhancers respond to distinct inputs, which means that the nature of the enhancer driving transcription dictates how a gene is regulated. There are many classic examples where these different enhancers are not on at the same time^2^. However, recent advances in single-cell sequencing and live imaging have revealed that multiple enhancers are often active when a cell is undergoing important developmental transitions^2,4,5^. Hence, carefully coordinating enhancer activity is vital for a gene to execute its function properly, particularly when enhancers that respond to disparate inputs are active simultaneously. How the activity of multiple enhancers is choregraphed in time remains poorly understood, mainly due to technical challenges in visualizing enhancer activity in real time. To understand how the action of different enhancers is coordinated *in vivo*, we examined a model system for specifying cell identity in the developing fruit fly (**Fig 1a**).

**Figure 1.**
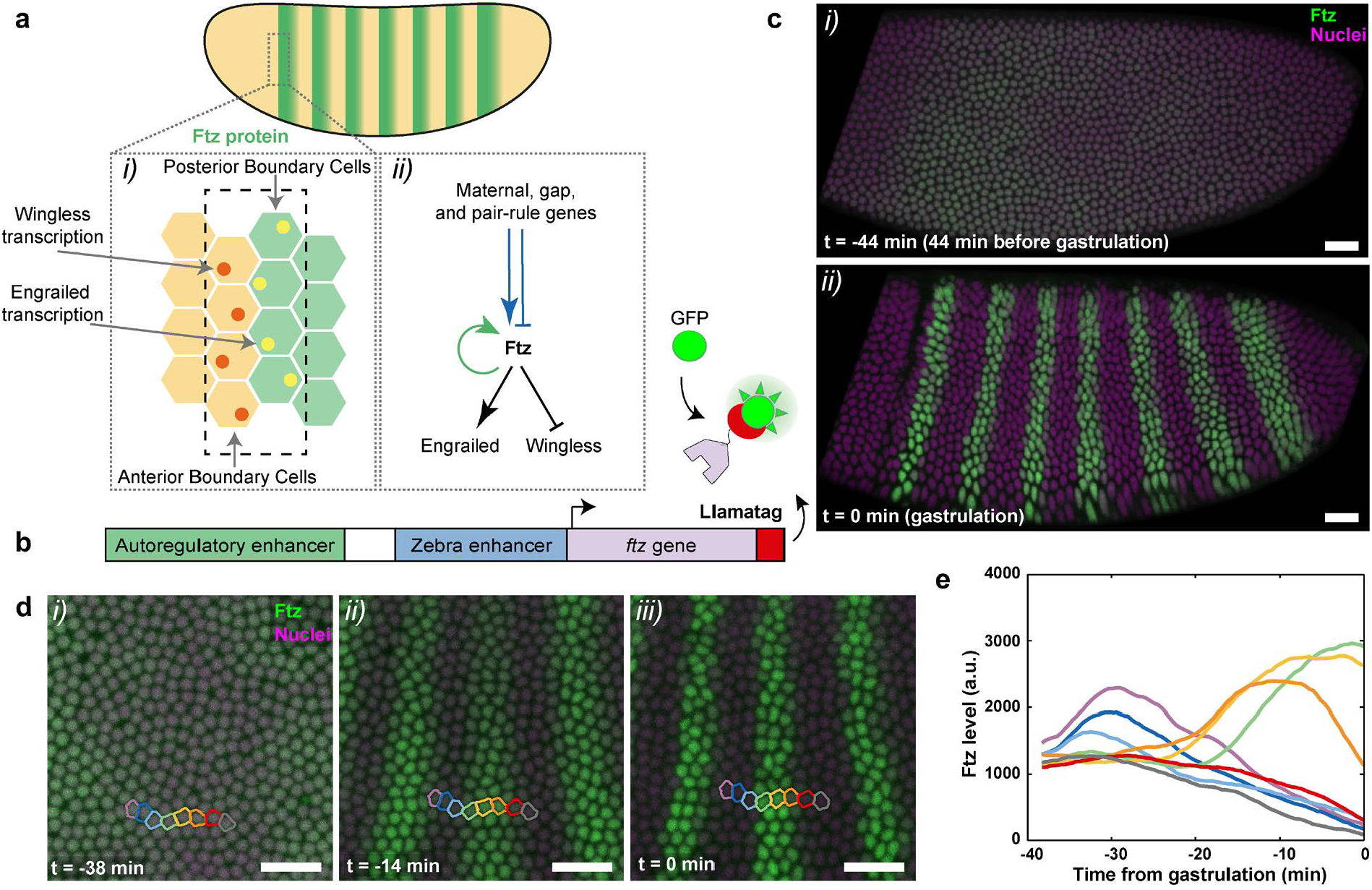
*Fushi tarazu* (*ftz*) regulatory network and pattern dynamics. **a**. Ftz forms a pattern of seven asymmetric stripes in the early fly embryo with the anterior edge defining compartment boundaries (*top*). *-i)* The anterior edge of each Ftz stripe consists of high-Ftz Posterior Boundary (PB) nuclei and low-Ftz Anterior Boundary (AB) nuclei. The Ftz target gene *engrailed* is expressed in the PB nuclei, while *wingless* is expressed in the AB nuclei. *-ii)* Maternal, gap and pair-rule genes regulate the transcription of *ftz*. Ftz directly activates *engrailed* and also its own transcription, while it indirectly inhibits *wingless*. **b**. Schematic of the endogenous *ftz* gene, which is regulated mainly by the autoregulatory and zebra enhancers, tagged with the LlamaTag. Maternally deposited eGFP binds the LlamaTag when present. **c**. Ftz pattern dynamics. -*i)* The early Ftz pattern, 44 minutes prior to gastrulation. -*ii)* The mature 7-striped Ftz pattern at gastrulation. **d**. Time course of the formation of Ftz stripes 3, 4 and 5. **e**. Time traces of Ftz levels for the color-coded nuclei in panel d. Scalebars are 25 µm.

The transcription factor Fushi tarazu (Ftz) plays an important role in patterning the early embryo where its expression is mainly controlled by two different enhancers: autoregulatory and zebra^6–10^ (**Fig 1b & Fig S1**). Over the course of an hour the initially broad Ftz pattern matures into a series of seven asymmetric stripes^11,12^. The anterior edges of these stripes specify the even-numbered compartment boundaries of the embryo^13–15^ (**Fig 1a-i**). These boundaries play a crucial role in development as signaling centers that are maintained throughout the life of the fly^16^. The anterior edges of the Ftz stripes are sharply defined, with a single row of nuclei containing high levels of Ftz positioned right next to a row of low-Ftz nuclei^17^. This binary protein pattern is essential for setting up the compartment boundary, by defining where two different selector genes, *engrailed* and *wingless*, are expressed^14,15,18,19^. How the two *ftz* enhancers work together to define the Ftz expression pattern is still unclear, but individually they have been extensively characterized.

It is known that both the autoregulatory and zebra enhancers drive expression in all seven Ftz stripes^7–9,20^. We also know that the zebra enhancer comes on first, but exactly when each enhancer is activated and repressed remains unclear. In general, the autoregulatory and zebra enhancers are regulated by various maternal, gap and pair-rule genes^7–9,20–22^ (**Fig 1a-ii**). However, in the rows of nuclei around the anterior edge of the stripes, *ftz* regulation is significantly simpler, being dominated by Ftz^23^. Here, Ftz protein is able to activate *ftz* transcription, and it is necessary to maintain transcription as the boundary sharpens^21,23^. The autoregulatory enhancer is thought to be responsible for this, because it contains multiple binding sites for Ftz that are necessary for the enhancer to switch on^8,9,20^. Despite all that is known, there are two key open questions about how the activity of these two different enhancers is coordinated to specify Ftz function.

What role does each enhancer play in shaping the anterior edges of the Ftz stripes? The prevailing view is that the zebra enhancer produces graded bands of Ftz protein, which are then thresholded by the autoregulatory enhancer to make an edge^8–10,21^. However, previous observations can also be explained by an opposing model. Here the zebra enhancer could be solely responsible for defining the sharp expression pattern, and the autoregulatory enhancer is activated later to simply maintain it. The second pressing question is: how exactly is the autoregulatory enhancer regulated? Is it switched on passively by the zebra enhancer, *via* the Ftz protein it produces, or is the timing of its initiation more directly regulated by other factors? While Ftz is clearly necessary to activate this enhancer^9^, it is also bound by other transcription factors^24^. If any of these are also necessary for activation and are expressed after Ftz, they would define the moment when the enhancer becomes active. Addressing these questions requires making precise real time measurements of gene activity in living embryos which was not possible until recently^25^.

## Results

### Boundary cells are identified at either side of the sharp anterior edges of Ftz stripes

To characterize the dynamics of the Ftz protein pattern, we fluorescently labeled the endogenous gene using the recently developed LlamaTag system^26^ (**Fig 1b**). Using this approach, we can accurately quantify protein dynamics in real time because the fluorescence we measure is directly proportional to protein concentration. This is not the case for standard fluorescent protein fusions where fluorescence is delayed and distorted by chromophore maturation effects, which is particularly problematic for short lived proteins like Ftz^26,27^. We then followed the Ftz pattern in living embryos using fluorescence confocal microcopy (**Fig 1c**). Multiple embryos were imaged for about an hour, and to reliably compare dynamics across individuals we measured time relative to when gastrulation occurred in each. The Ftz pattern evolved from a broad domain of expression to seven distinct stripes, consistent with previous observations^26,28^ (**Fig 1c & Supplemental movie 1**). In addition to following the pattern as a whole we could also track individual nuclei (**Fig 1d**), and reliably quantify how the concentration of Ftz protein changed in each nucleus over time (**Fig 1e**). Using this robust system we focused on the nuclei that define the compartment boundary to understand how they assume different fates.

In order to unambiguously identify the nuclei at the anterior edge of each Ftz stripe, we examined the refined pattern just before gastrulation, and classified these high-Ftz nuclei based on the Ftz levels at this point (**Fig 2a & Fig S2**). Similarly, we could identify the row of nuclei, with low Ftz levels, just to the anterior of these. We refer to the high-Ftz border row of cells as the Posterior Boundary (PB) nuclei, since they are posterior of the *compartment* boundary, and, similarly, the row of neighboring low-Ftz nuclei as the Anterior Boundary (AB) nuclei (**Fig 2a-ii**). After classifying the nuclei at either side of the compartment boundary into either low-Ftz (AB) or high-Ftz (PB) categories, we traced the Ftz level trajectories of these nuclei back in time to see how they arrived at these different final states (**Fig 2b**).

**Figure 2.**
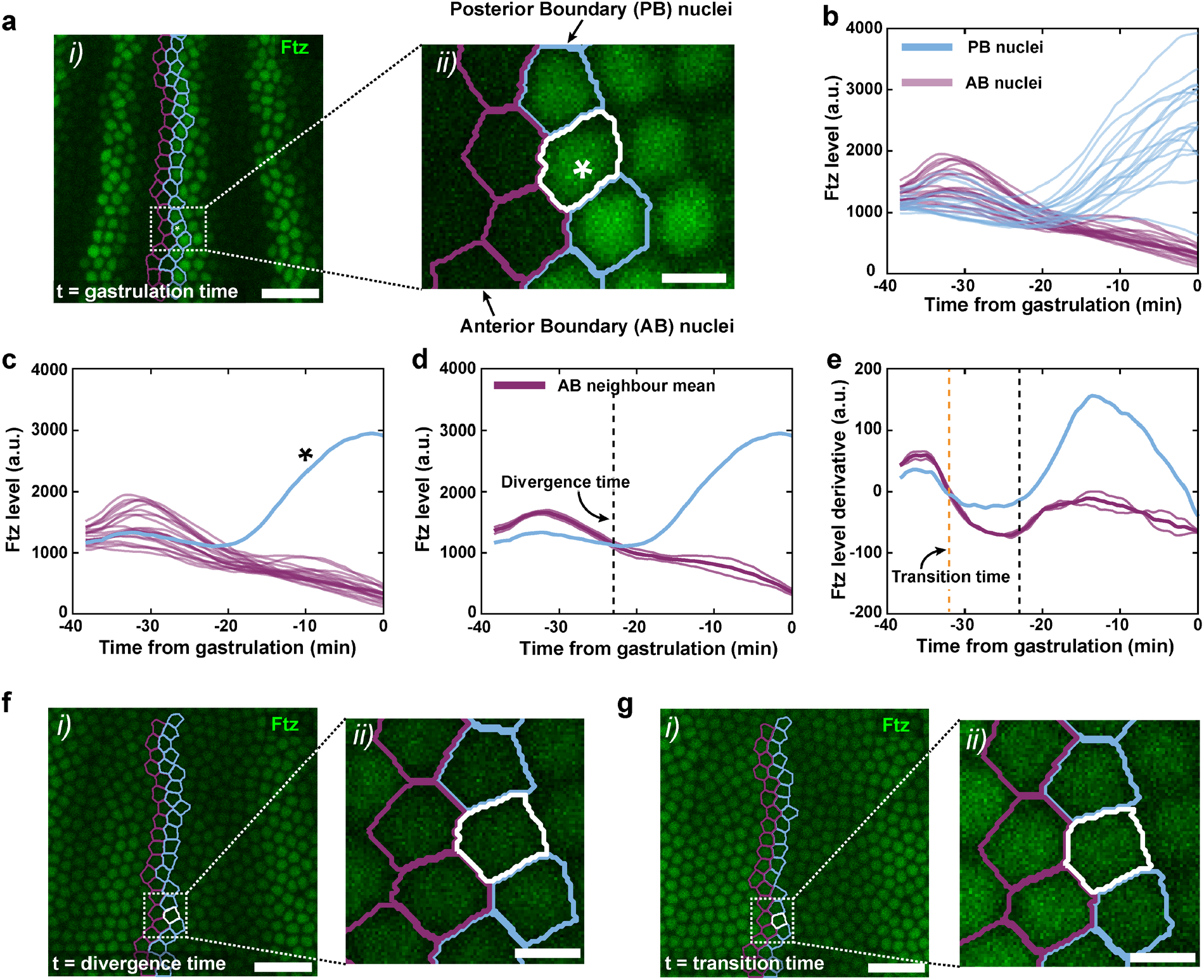
Transition time as an estimate for when boundary nuclei make their fate choice. **a***-i)* Ftz stripes 3, 4, and 5 at gastrulation. *-ii)* zoom-in of *i)*. PB nuclei are outlined in blue and AB nuclei are shown in purple. Asterisk indicates the PB nucleus used in panels c, d, and e (see also **Fig S2**). **b**. Ftz traces of PB and AB nuclei in stripe 4 of a single embryo. **c**. Ftz time traces of the PB nucleus highlighted with an asterisk in panel a, together with all AB nuclei. **d**. Ftz time trace of the PB nucleus highlighted with an asterisk in panel a, together with the traces of its two AB neighbors, and their mean (thick purple line). The divergence time is annotated by a black dashed line. **e**. Ftz level derivative of the equivalent nuclei in **d. f** and **g**. The Ftz pattern at the divergence (**f***-i)*, and the transition time (**g***-i)* of the PB nucleus outlined in white. *-ii)* zoom-in of *i)*. Scalebar is 25 µm for overview panels and 5 µm for the zoom-in panels.

### The transition time estimates the timepoint at which boundary nuclei commit to their cell fate

The AB and PB nuclei are clearly different at gastrulation, but only 40 minutes earlier they are indistinguishable based on Ftz levels alone (**Fig 1c**). Using the Ftz time traces and knowledge of the final state, we sought to determine the earliest point in time when we could deduce if a nucleus would finish in the high or low Ftz state. To ensure that our approach was robust to any systematic differences that may exist in nuclei at different positions, we only compared the traces of individual PB nuclei to the average of their nearest AB neighbors (**Fig 2c & 2d**). Given that at gastrulation a PB nucleus ends up with high Ftz levels, and the neighboring AB nuclei with low Ftz levels, there must exist a *latest* point at which the Ftz level trajectories intersect. We defined this timepoint, at which the Ftz protein traces of a PB nucleus and its AB neighbors intersected for the last time, as the “divergence time”. After this divergence time, the Ftz trace of a PB nucleus continues its upward trajectory, while the Ftz levels of its AB neighbors continue to decrease.

Upon careful inspection of many protein traces, we noted differences in the trajectories of the AB and PB nuclei even before the divergence time. This makes sense if one considers that a PB nucleus must increase Ftz production at an earlier time in order to set out on a consistent upward trajectory after the divergence time. Similarly, the neighboring AB nuclei must embark on a trajectory of net protein degradation, in order to consistently decrease their Ftz levels after the divergence time. This implies that there exists a timepoint *before* the divergence time, at which the net protein production/degradation in a PB and its AB neighbors were equal for the last time. The balance between protein production and degradation is proportional to the rate of change of the Ftz levels, *i*.*e*. the derivative of the Ftz traces. By following the Ftz level derivatives of a PB nucleus and its AB neighbors backwards in time from the divergence point, we determined the *latest* time at which these derivatives intersected. We defined this timepoint as the “transition time”, and we observed that it generally preceded the divergence time by about 10 minutes (**Fig 2e**). The dynamic protein data provides a much clearer view of how these nuclei make developmental decisions compared to what can be learned from snapshots of fixed samples, as is typically done. This is made particularly obvious by looking at snapshots of the Ftz pattern at the divergence and transition times (**Fig 2f & 2g**). Based on these pictures alone, it would not be possible to identify which nuclei would eventually express high or low levels of Ftz. Our approach, on the other hand, allows us to identify a thus far undescribed timepoint at which cell identity is acquired.

### Boundary nuclei of different stripes make their fate decision at different times

To determine how consistent the transition time was across multiple embryos and between different stripes, we examined 3 stripes in 10 to 12 embryos and determined the transition times for 455 nuclei (**Fig 3a & 3b**). The median transition times for stripe 3 and stripe 4 were relatively similar at -27 min and -25 min, respectively, with stripe 5 occurring earlier at -33 minutes. The earlier average transition time for stripe 5 is consistent with results from previous studies showing it forms before stripe 3 and 4^10,28^. To establish if the variation in the transition time values within a stripe was partly due to a systematic spatial bias, we calculated the difference in the transition times of neighboring nuclei and plotted that distribution for the three stripes (**Fig 3c**). The distribution for all three stripes is similar and sharply peaks at 0 minutes, indicating that much but not all of the spread seen in **Fig 2b** comes from nuclei in different dorsal-ventral positions being slightly offset from one another in time. We conclude that each stripe has its own characteristic range of transition times, with spatially proximal nuclei being more similar.

**Figure 3.**
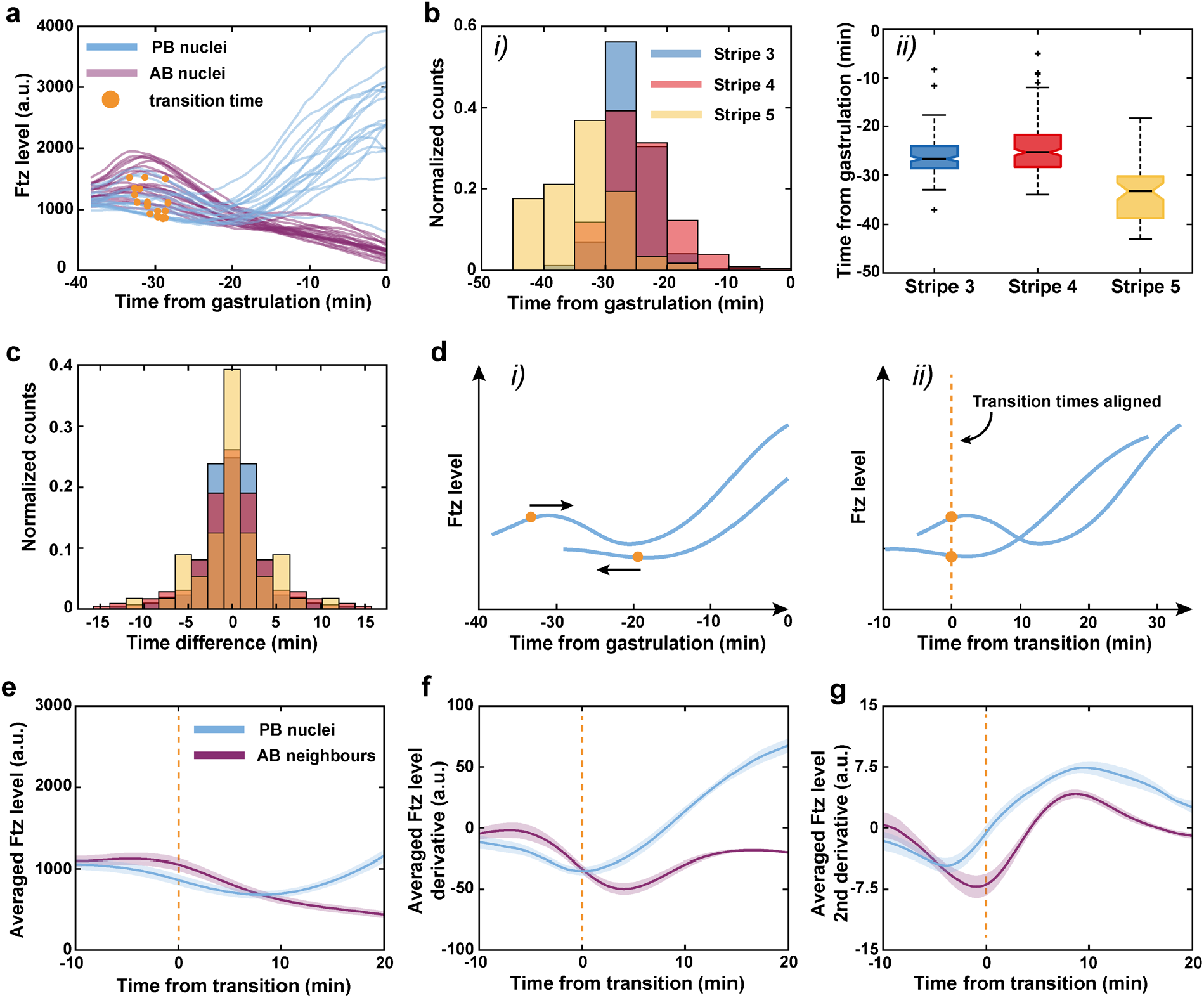
Characterization of the transition times and re-alignment of Ftz traces. **a**. Ftz traces in stripe 4 of a single embryo with the transition times denoted by orange dots. **b**. The transition time relative to gastrulation for stripes 3 (171 nuclei in 10 embryos), 4 (227 nuclei in 12 embryos), and 5 (57 nuclei in 10 embryos) displayed as *-i)* a histogram and *-ii)* a boxplot. **c**) Differences in transition times between neighboring PB nuclei. **d**. Schematic illustrating the realignment of Ftz traces relative to their respective transition times. *-i)* Ftz level traces before realignment. *-ii)* Ftz time traces are shifted until the transition times are re-aligned at t = 0. **e -g** Averaged Ftz levels, 1^st^ derivative and 2^nd^ derivative relative to the transition time of PB nuclei and their AB neighbors for all stripe 4 nuclei. In panels e, f, and g, the thick line shows the mean and the shading the Standard Error of the Mean, due to variations across embryos (see **Fig S3** for stripe 3 and 5).

### The 2^nd^ derivatives of the protein traces predict differential transcription in boundary nuclei before the transition time

We next wanted to learn more about how the AB and PB nuclei were making their fate decision by looking for systematic trends in their Ftz dynamics. These trends might not be obvious from single traces, but could be more readily observed in the averages of many traces. However, since all nuclei have different transition times, the developmental trajectories are also slightly shifted relative to each other. This means that taking a simple time average of all the traces would obscure features around this important timepoint. To get around this problem, we chose to re-align the traces in time to their respective transition times, as shown by the schematic in **Fig 3d**. By comparing the averages of the aligned Ftz traces for the PB nuclei and their AB neighbors (**Fig 3e & Fig S3a/d**), we could see the two classes of nuclei had equal Ftz levels about 8 minutes after the transition time, which is consistent with our earlier observation that the transition time preceded the divergence time by this amount. Similarly, we could verify that the averaged first derivative of Ftz levels in PB nuclei and their AB neighbors intersected at the transition time, as per its definition for individual traces (**Fig 3f** and **Fig S3b/e**).

The surprising result came when we looked at the second derivative of the Ftz traces (**Fig 3g & Fig S3c/f**), which is closely related to how transcription is regulated^29^. Indeed, as long as there are no time-dependent changes in translation, the second derivative of a protein trace is directly proportional to changes in its mRNA levels^29^. These changes in turn are dominated by the rate at which the gene is being transcribed, with a characteristic delay related to how long it takes mRNA to be synthesized and exported from the nucleus. As a result, the first indication that there is a change in the transcription of a gene will appear as a change in the second derivative of its protein traces. In our comparison of the PB nuclei and their AB neighbors, we can clearly see their second derivative traces intersect around 5 minutes before the transition time. This indicates that around this time Ftz starts to be differentially transcribed in the two classes of nuclei. Taking into account the time it takes to transcribe and export the mRNA (∼3 minutes^23,30^), we expect there to be a change in how *ftz* is transcribed in PB and AB nuclei about 8 minutes before the transition time.

### The autoregulatory enhancer activates long after the transition time

To understand how the autoregulatory and zebra enhancers regulate this Ftz fate decision we measured their activity over time. We started by creating a transcriptional reporter of the autoregulatory enhancer using a large (2.6 kbp) fragment containing the enhancer, as well as the endogenous *ftz* transcription unit with the native promoter (**Fig 4a**). This reporter also had 24 copies of the MS2 stem loop sequence (which binds to MCP-mCherry) inserted into the intron to visualize transcription live^31–33^. We could then measure the activity of the autoregulatory enhancer (as dots in the mCherry channel) while simultaneously following changes in Ftz protein levels in the same nuclei (**Fig 4b-d)**. This reporter drove strong expression in seven stripes, consistent with what had been reported previously from fixed samples^9,20^ (**Supplemental movies 2 & 3**), but had some unexpected dynamic properties.

**Figure 4.**
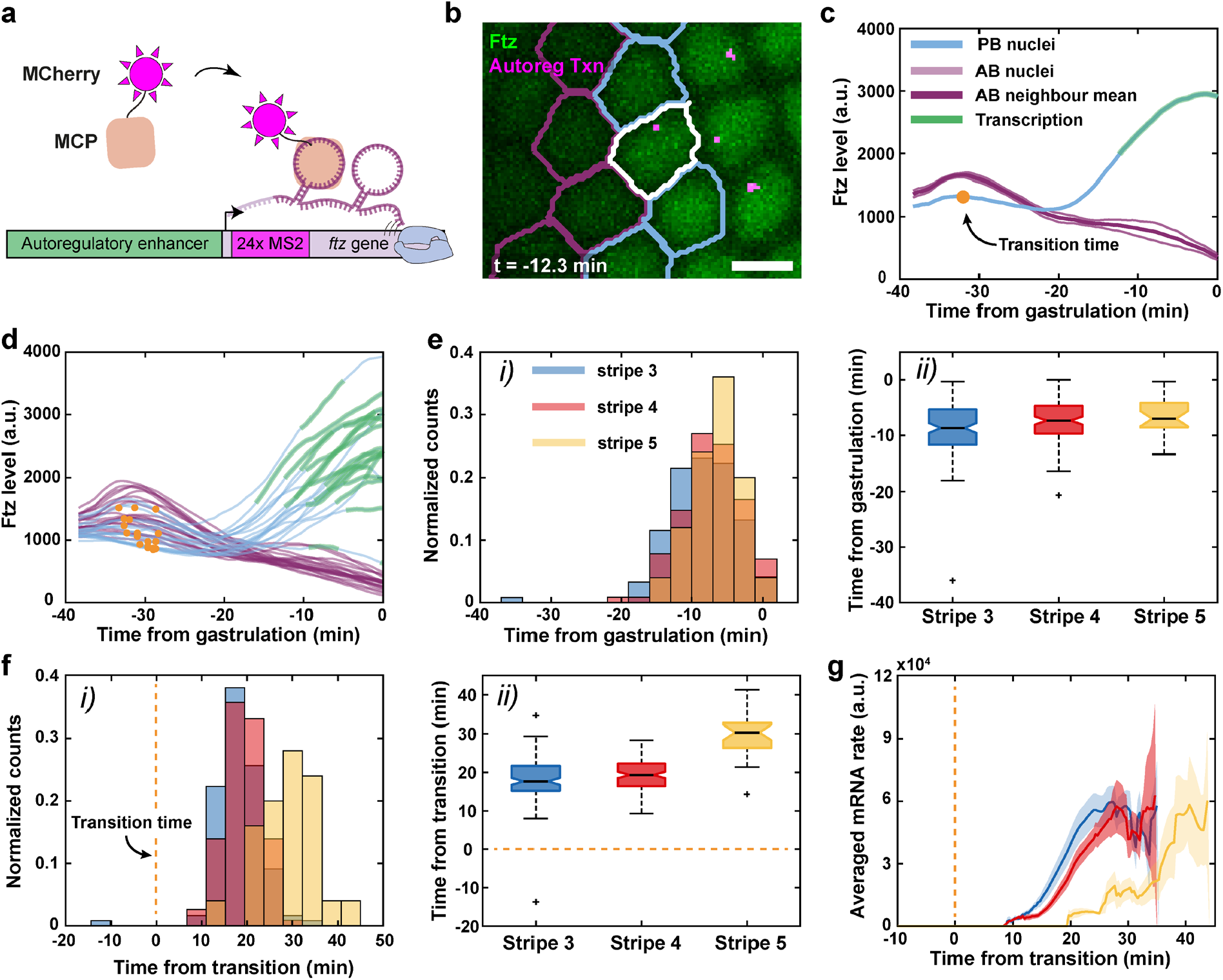
Autoregulatory enhancer activates transcription after the transition time. **a**. Schematic of MS2 based approach to visualize transcription of autoregulatory enhancer in real time. **b**. The Ftz pattern at the timepoint of the first mRNA spot in the PB nucleus outlined in white. **c**. Ftz level traces of the PB nucleus outlined in white from panel b, together with its two AB neighbours and their mean. The transition time is annotated with an orange dot, and the time period during which transcription is observed is in green. **d**. Ftz level traces of PB nuclei showing transcription, and all the AB nuclei. **e**. The first activation times of the autoregulatory enhancer relative to the gastrulation time in PB nuclei across stripe 3, 4, and 5, displayed as -*i*) a histogram; and -*ii*) a boxplot. (see also **Fig S4**) **f**. Same as panel e, but with the first activation times re-aligned relative to the transition time of the PB nuclei, displayed as -*i*) a histogram; and -*ii*) a boxplot. **g**. Averaged mRNA rate relative to the transition time in stripes 3, 4, and 5. The thick line shows the mean and the shading the Standard Error of the Mean, due to variations across embryos. Nuclei and embryo numbers used for panels e and f: stripe 3 -110 nuclei in 10 embryos; stripe 4 - 115 nuclei in 12 embryos; stripe 5 - 25 nuclei in 10 embryos. Scale bar is 5 µm.

Surprisingly, the autoregulatory enhancer became active long after the transition time, contrary to the dominant role it was expected to play in shaping the cell fate decisions of PB and AB nuclei. For stripe 4, we only see sustained bursts of transcription about 19 minutes after the transition time, at which stage the Ftz levels in AB and PB nuclei are already very different (**Fig 4d**). Moreover, the enhancer became active at similar times in stripe 3, 4, and 5 (between -7 and -9 min), despite these individual stripes forming at different times^12,28^ (**Fig 4e**). The trend of late activation of the autoregulatory enhancer, up to 30 minutes after the transition time, was consistent in the data across many different nuclei, embryos, and the three different stripes (**Fig 4f & 4g**). This left us to conclude that the autoregulatory enhancer cannot be responsible for driving the fate decision of the AB and PB nuclei because it is activated too late.

### Differential transcription driven by the zebra enhancer in boundary nuclei precedes the transition time

Next, we sought to establish if the zebra enhancer was responsible for defining the edges of the Ftz pattern. Similar to the autoregulatory enhancer, we created a transcriptional reporter for the zebra enhancer (**Fig 5a**). We observed that the zebra enhancer starts to drive transcription much earlier than the autoregulatory enhancer, and is initially detected broadly across the embryo. It then becomes progressively restricted to seven stripes, and finally dissipates around gastrulation (**Supplemental movies 4 & 5**). In order to compare the activity of the zebra enhancer and the timing of the endogenous Ftz protein dynamics, we chose to focus on stripe 4, because it has no known “shadow” stripe enhancer^21^ (**Fig S1**). These enhancers usually serve to improve patterning robustness in response to stress, but may also influence Ftz concentration kinetics. Around 40 minutes before gastrulation, the average rate at which the zebra enhancer drove transcription was similar in the PB nuclei and their AB neighbors, but by gastrulation they were very different (**Fig 5c**). Interestingly, the PB nuclei and AB neighbors started to show clear differences in zebra transcription rate right around their transition times.

**Figure 5.**
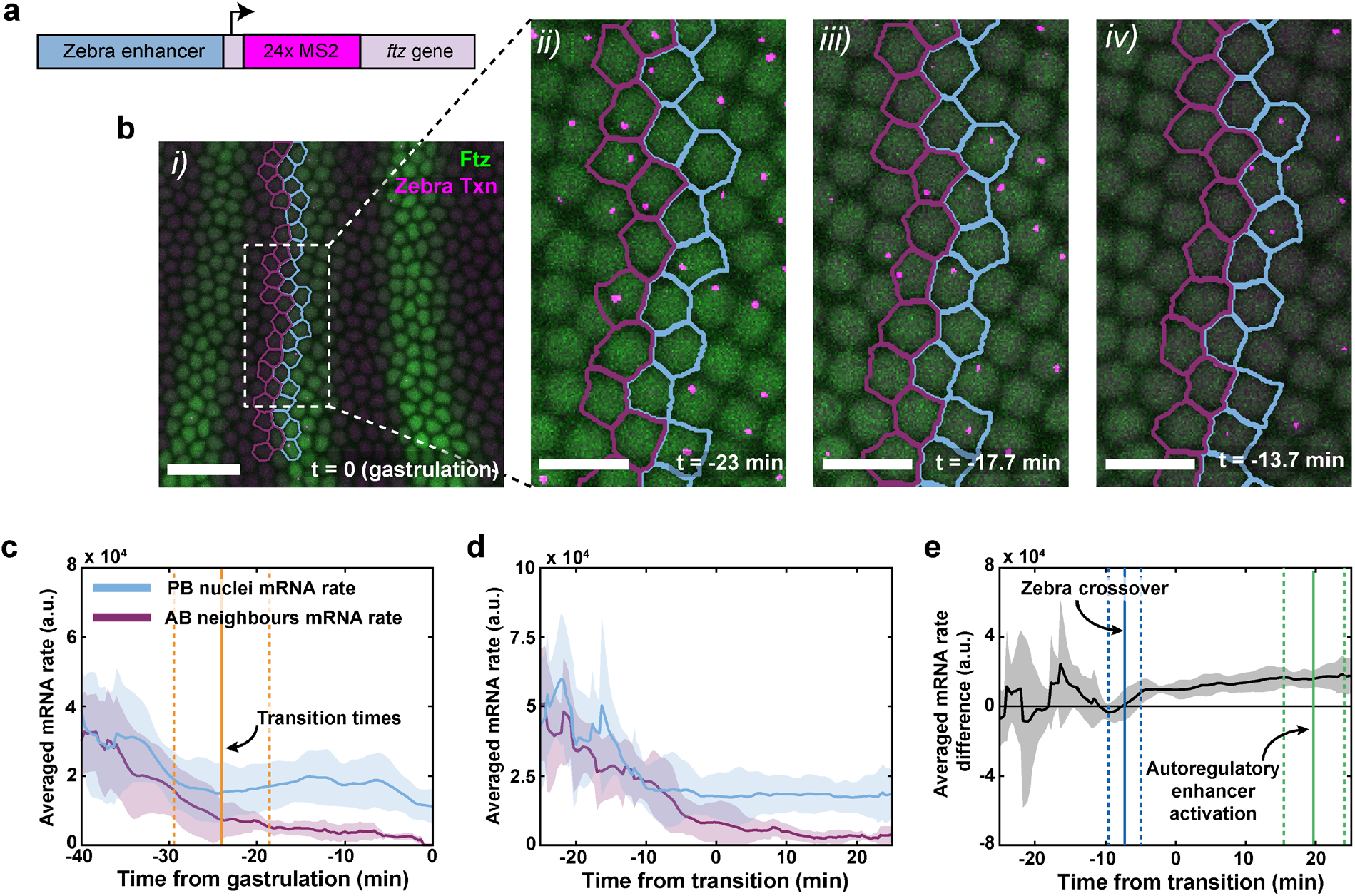
Zebra enhancer is differentially transcribed at the boundary before the transition time. **a**. Schematic of MS2 based approach to visualize transcription of the zebra enhancer in real time. **b**-*i*) Ftz pattern and zebra transcription in stripe 3, 4 and 5 at gastrulation time, with AB and PB nuclei in stripe 4 annotated. Zoom-ins are shown at different times (*ii* to *iv*) Note how zebra transcription disappears from the AB nuclei, while it stays present in PB nuclei. **c**. Averaged mRNA production rate in the PB nuclei of stripe 4 and their AB neighbors, relative to gastrulation time (thick line is the mean and the shading the standard deviation). The range of transition times are annotated by the thick orange line (mean across stripe 4) and dashed lines (standard deviation). **d**. The averaged mRNA rate in stripe 4 PB nuclei and their AB neighbors, relative to the transition times of the PB nuclei (thick line is the mean and the shading the standard deviation). **e**. The averaged difference in mRNA rate between stripe 4 PB nuclei and their AB neighbors. The range of activation times of the autoregulatory enhancer in stripe 4 is annotated with a thick green line (mean) and dashed lines (standard deviation). The range of times at which the transcription rate in stripe 4 PB nuclei consistently outgrows that in AB neighbours is manually annotated with a thick blue line (mean mRNA rate difference crosses zero for last time) and dashed lines (the standard error on the mean crosses zero). Scalebar is 25 µm for overview panels and 5 µm for the zoom-in panels. See **Fig S5** for stripe 3 and 5.

To clarify trends around the transition times, we re-aligned the transcription traces of each PB nucleus and its AB neighbors to the respective PB transition time (**Fig 5d**). At the transition time, the zebra enhancer already drives transcription at a higher rate in the PB nuclei than in their AB neighbors. The time at which they first start to show differential transcription (*i*.*e*., the last time at which the transcription rates are equal in the two nuclei classes) occurs about 8 minutes before the transition time (**Fig 5d & 5e**). This coincides with when we expected a change in how *ftz* is transcribed in PB and AB nuclei based on the second derivatives of their Ftz levels (**Fig 3g**). This strongly suggests a connection exists between the cell-fate decision obtained from features in the protein traces and differences in zebra-driven transcription. Taken together, this supports the idea that the zebra enhancer is the enhancer responsible for defining the sharp anterior edge of the Ftz stripe.

### The autoregulatory and *engrailed* enhancers are time-controlled

If the autoregulatory enhancer is not defining the edge, then it likely serves to maintain the decision defined by zebra. To address how this is regulated we needed to understand precisely how the autoregulatory enhancer is responding to Ftz. For each nucleus we determined at what Ftz concentration the autoregulatory enhancer was first activated (Ftz_a_), and at what time this activation occurred (t_a_) (**Fig 6d**). Using this data, we tested if the autoregulatory enhancer switched on at a particular Ftz concentration (threshold model), or if instead it was activated after a specific time set by some other factor (timer model) (**Fig 6b & 6c**). To this end, we compared the distributions of observed Ftz_a_ and t_a_ values to control distributions of Ftz levels and activation times generated from the Ftz traces under assumption of either the threshold or timer model (see STAR-methods). We then asked if the observed and control distributions were statistically different (**Fig 6e**). Our results show that the t_a_ distribution is different from the control distribution, while the same is not true for the Ftz_a_ distribution. We saw this for each of the stripes used in these experiments (**Fig S6a-d**), which taken together strongly support a timer model over a threshold model for the activation of the autoregulatory enhancer.

**Figure 6.**
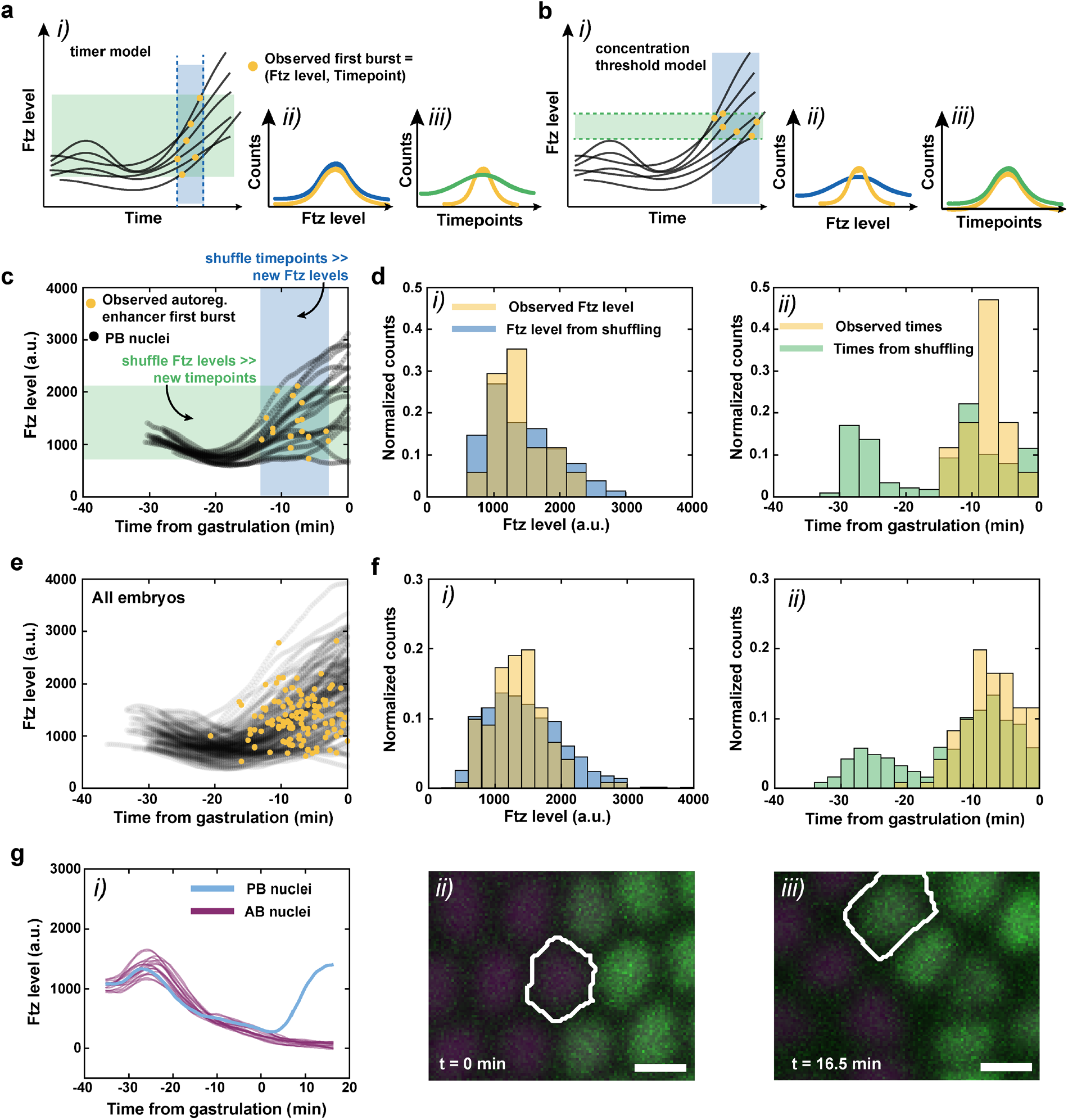
The autoregulatory enhancer is activated at a characteristic time, not when Ftz reaches a specific level. **a**-*i*) In a “timer model”, transcription (yellow dots) initiates during a defined time window (blue shading and dashed lines) and across a range of Ftz levels which correspond to those present within the time window (green shading). -*ii*) If the observed timepoints are shuffled, and one searches for the new associated Ftz levels (blue line) within the blue shading of panel a-*i*, one recovers the same Ftz level distribution as observed (yellow line). -*iii*) If the observed Ftz levels are shuffled, and one searches for the new associated timepoints (green line) within the green shading of panel a-*i*, one recovers a broader distribution than observed (yellow line). **b**-*i*) In a “threshold model”, transcription (yellow dots) occurs at defined Ftz levels (green shading and dashed lines) and across a range of timepoints corresponding to the times at which those Ftz levels occur (blue shading). - *ii*) If the observed timepoints are shuffled, and one searches for the new associated Ftz levels (blue line) within the blue shading of panel b-*i*, one recovers a broader Ftz distribution than observed (yellow line). -*iii*) If the observed Ftz levels are shuffled, and one searches for the new associated timepoints (green line) within the green shading of panel b-*i*, one recovers the same timepoints distribution as observed (yellow line). **c**. Scatter plot of Ftz levels in nuclei displaying transcription (stripe 4, a single embryo). **d**-*i)* Observed Ftz levels at first autoregulatory enhancer transcription burst (yellow) and Ftz levels (blue) obtained from the shuffling procedure described in panel a/b-*ii. -ii)* Observed times of first autoregulatory enhancer transcription burst (yellow) and timepoints (green) obtained from the shuffling procedure described in panel a/b*-iii*. **e** and **f**. Same as panels c and d, but applied to all 12 embryos with stripe 4. **g**-*i*) Examples of a PB nucleus in stripe 3 of a zebra enhancer embryo whose Ftz levels increase after gastrulation time. For comparison, the traces of AB nuclei are displayed too. -*ii* and -*iii*) Images of the PB nucleus (white outline) at gastrulation time (-*ii*) and at a later time when the Ftz levels have increased (-*iii*). Two-sample Kolmogorov-Smirnov test at 5% significance was used for panels d and e. Null-hypothesis = observed and shuffled values come from same distribution. Alternative hypothesis = shuffled values tend to be smaller than observed values. For panel d-*i* and e-*i*, the test does not reject the null hypothesis. For panel d-*ii* and e-*ii*, the test does reject the null-hypothesis and accepts the alternative hypothesis. Scale bar is 5 µm. See **Fig S6** for stripe 3 and 5.

One of the key differences between the timer and threshold model is how they predict the autoregulatory enhancer will respond to low levels of Ftz. According to the timer model, even nuclei with very low levels of Ftz can induce the autoregulatory enhancer after the activation time and upregulate Ftz. The threshold model, on the other hand, predicts that nuclei with Ftz levels significantly below the threshold simply cannot induce the autoregulatory enhancer. The timer model would then predict that after the activation time, we should occasionally be able to find nuclei with low Ftz levels that become Ftz positive. We do indeed find these nuclei at low frequency in our data, which provides further support for the timer model (**Fig 6f & 6g**).

In order to determine if the timer model of Ftz-mediated activation could apply to other Ftz targets, we turned our attention to *engrailed*. The gene *engrailed* is a direct target of Ftz^23^ and to image its activation we created a PP7 tagged version of the endogenous gene using CRISPR^28,34^ (**Fig 7a & 7b**). Our tagged version of *engrailed* faithfully recapitulated endogenous expression^35,36^ (**Fig 7b**). The first *engrailed* transcription was observed around gastrulation, but most nuclei do not show expression until afterwards, by which time they have moved significantly. By correcting for signal intensity differences in nuclei at different depths, we were able to reliably perform quantitative analyses on the Ftz traces even 19 minutes after gastrulation. Performing the same analysis as was done for the autoregulatory enhancer, we found that the distribution of t_a_ values for *engrailed* was different from the control distribution, while that of Ftz_a_ was not (**Fig 7d/e & Fig S6e-h**), similar to the result of the autoregulatory enhancer. This provides strong support for the timer model over the threshold model for a second direct Ftz target, suggesting that this might be a more general way in which Ftz activates its targets. The observation is also consistent with earlier work showing that the timing of *engrailed* activation did not change as the copy number of Ftz was altered^14^. Interestingly, *engrailed* is activated about 10 minutes after the autoregulatory enhancer (**Fig 7c**), indicating that the precise value of the activation time could be enhancer specific.

**Figure 7.**
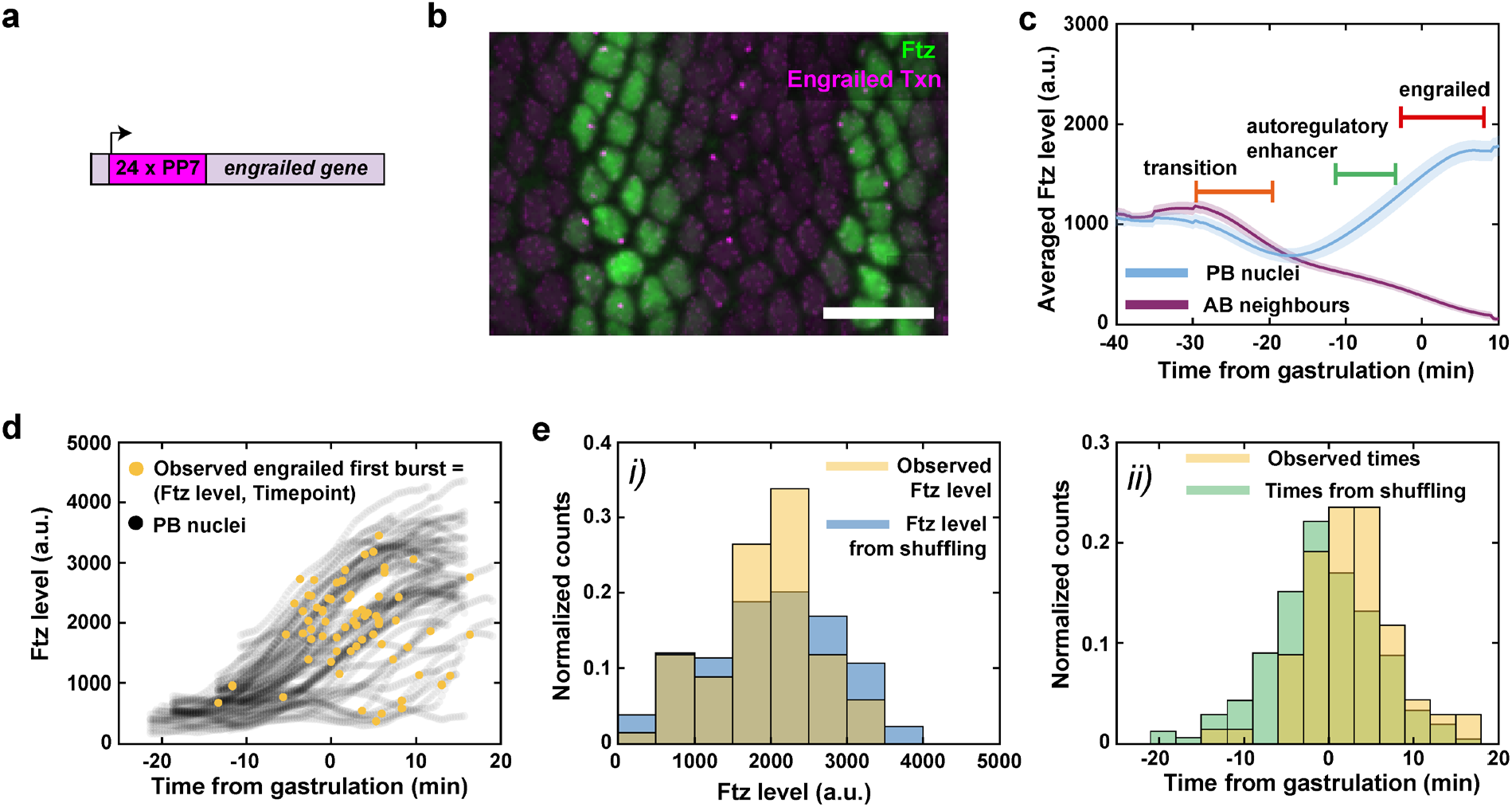
*Engrailed* transcription initiates at a characteristic time, not when Ftz reaches a specific level. **a**. Transcription of *engrailed* is visualized by the insertion of 24 PP7 repeats into the intron of the endogenous locus, and the subsequent binding of PCP-Tomato to the RNA stem loops after transcription has initiated. **b**. Image of the Ftz pattern after gastrulation together with the *engrailed* stripe mRNA dots driven by Ftz and also Eve. **c**. The averaged Ftz level (thick line is mean and shaded area is SEM) in PB nuclei and their AB neighbours in stripe 4 of autoregulatory enhancer embryos, together with the annotation of the time range (+/-standard deviation from the mean) of transition times, autoregulatory enhancer activation and engrailed activation in PB nuclei. **d**. Scatter plot of Ftz levels after the transition time in nuclei displaying transcription (stripe 4 in 5 embryos). **e.-***i*) Observed Ftz levels at first *engrailed* transcription burst (yellow) and Ftz levels (blue) obtained from the shuffling procedure described in panel a/b-*ii* of figure 6. *-ii)* Observed times of first *engrailed* transcription burst (yellow) and timepoints (green) obtained from the shuffling procedure described in panel a/b-*iii* of figure 6. For panel e-*i*, the Two-sample Kolmogorov-Smirnov test does not reject the null hypothesis, while for panel e-*ii* it does. Scale bar is 20 µm. See **Fig S6** for stripes 3 and 5.

## Discussion

We find that the zebra enhancer defines the anterior edge of stripe 4 early and precisely. This occurs by maintaining *ftz* transcription in the PB nuclei while it is systematically lost in the AB nuclei (**Fig 5**). Based on what is known about Ftz regulation, this decrease in transcription is most likely mediated by transcriptional repressors such as Even-skipped^37^. We also see no reason why a similar mechanism could not hold true for the other Ftz stripes, but then mediated by a combination of the zebra enhancer and their respective shadow stripe enhancers. Modelling how repressors specify the edge of Ftz stripes may provide a unique opportunity to extend the well-developed theoretical frameworks that connect enhancer sequence to boundary specification by activator proteins. The best-known examples of such studies are predictions for how the Bicoid activator gradient specifies the Hunchback boundary^38–42^. Compared to Hunchback, the Ftz boundaries are specified much faster and are much sharper, which will pose new challenges to existing models that would likely yield fundamentally new insights.

Our results argue that the autoregulatory enhancer “locks in” the Ftz fate decision defined by the zebra enhancer, rather than refining the Ftz protein pattern initially produced by zebra. This ensures that Ftz expression is maintained even after the zebra enhancer is no longer active. The ability of the autoregulatory enhancer to respond to even low levels of Ftz protein ensures that it can rapidly “lock in” the Ftz positive state from a broad range of starting Ftz concentrations. However, it also means the autoregulatory enhancer needs to be kept inactive while the Ftz pattern evolves from its initially broad domain of expression to well-defined stripes. So, it is important to regulate precisely when the autoregulatory enhancer is activated, and recent work has uncovered possible mechanisms by which this can occur.

It has been shown that the ubiquitously expressed transcription factors Caudal, Dichaete and Odd-paired behave like simple timers within the early embryo^22,43–45^. These three factors are sequentially expressed, bind to a diverse range of enhancers, and have pioneer-like activity that modulate chromatin state. In fact, Odd-paired has been shown to open the chromatin of a subset of enhancers that include the Ftz-dependent enhancers of *engrailed*^44^. ChiP data also shows that the autoregulatory enhancer is bound by both Dichaete and Odd-paired^46^. We hypothesize that these global timing factors control when the autoregulatory enhancer’s chromatin is opened, after which it can respond to Ftz. The difference in activation timing between the autoregulatory enhancer and *engrailed* could be caused by predominant activation of *engrailed* through Odd-paired, which peaks later than Dichaete. Additionally, the timing difference might be related to differential affinity of the Odd-paired binding sites in the autoregulatory and engrailed enhancers. The concentration of Odd-paired peaks at gastrulation, but during the preceding half-hour window its concentration monotonically increases^44^. Hence, the affinity of Odd-paired binding sites could define exactly when an enhancer becomes active during this time.

The precise control of when the zebra and autoregulatory enhancers are active is necessary to properly define the anterior edges of the Ftz stripe pattern, and by extension, the compartment boundaries. Furthermore, during development many processes, such as cell-fate specification and morphogenesis, are happening in parallel and need to remain coordinated. We speculate that the aforementioned global timing factors, which regulate the relative activation times of various enhancers, provide a means for ensuring this coordination. Having global timing factors regulate the transitions between different enhancers across the range of genes that drive these processes provides a simple mechanism to ensure that building an embryo runs like clockwork.

## Acknowledgments

We thank Matthew Norstad for early technical assistance with the preparation of genetic constructs, Anika van der Zant for help with imaging and Emilia Cadar, Koen Schuddemat, Cemil Korkmaz, and Jochem Boeter for advice on experiments and analysis. We gratefully acknowledge Catherine Rabouille, Katharina Sonnen, Juan Garaycoechea and Francesca Mattiroli for insightful feedback on the manuscript. We also thank the members of the Bothma, Sonnen, and Garaycoechea labs for fruitful discussions and thank the members of the Hubrecht Media Kitchen and Imaging Facility for experimental support. This study was financially supported by the Hubrecht Insitute and ERC starting grant (TF-Dynamics-949708).

## Author contributions

Conceptualization, J.P.B., A.B., A.P.; Methodology, A.P. and J.P.B.; Software. A.B. and J.P.B.; Formal Analysis, A.B.; Investigation, A.P.; Data Curation, A.B.; Writing - Original Draft, J.P.B., A.B., A.P.; Writing - Review & Editing, J.P.B., A.B., A.P.; Visualization, J.P.B., A.B., A.P.; Supervision, J.P.B.; Project Administration, J.P.B.; Funding Acquisition, J.P.B.

### Declaration of interests

The authors declare no competing interests.

## STAR Methods

### RESOURCE AVAILABILITY

#### Lead contact

Further information and requests for resources and reagents should be directed to and fulfilled by the lead contact, Jacques P. Bothma.

#### Materials availability

- All plasmids produced and fly lines generated in this paper will be shared by the lead contact upon request.

#### Data and code availability

- All imaging data reported in this paper will be shared by the lead contact upon request.
- All original code has been deposited <u>here</u> and is publicly available as of the date of publication. DOIs are listed in the key resources table.
- Any additional information required to re-analyze the data reported in this paper is available from the lead contact upon request.

## EXPERIMENTAL MODEL AND SUBJECT DETAILS

The experimental model used in this study is *Drosophila melanogaster*. All individuals used in this study were embryos that were imaged as detailed below during the first 5 hours of development. Embryos were allowed to develop at room temperature and conditions unless otherwise stated. Embryo sex is not reported as it is not believed to influence any of the measurements reported here.

### Fly Strains/Genotypes

The unpublished fly lines that were used in this study were generated by incorporating engineered transgenes into the genome of the *yw* fly strain, or by altering endogenous loci of the *yw* strain using CRISPR-Cas9 mediated homologous recombination. The Cloning and Transgenesis section details how each transgene was generated and genomically integrated, as well as how specific loci in the genome were edited using CRISPR-Cas9 mediated homologous recombination.

#### Fushi tarazu Protein and MCP-mCherry-NLS

To image the expression of Ftz protein in the early embryo, we performed fly crosses to combine the integrated transgene that encoded maternal eGFP, and maternal MCP-mCherry-NLS which served as a nuclear marker. The full genotype of these mothers was *yw; vasa>eGFP; nos>*MCP-mCherry-NLS. Virgin females of this line were collected, crossed in a cage with male flies that were homozygous for the Ftz locus and that had been tagged with the eGFP LlamaTag (*yw;+; Ftz-LlamaTag)*. The resulting embryos were imaged. This ensured that the imaged embryos contained maternally deposited eGFP, MCP-mCherry-NLS, and the tagged Ftz locus.

#### Fushi tarazu Protein and AutoregulatoryEnhancer-Ftz-MS2

To simultaneously image Ftz protein and transcription from the AutoregulatoryEnhancer-Ftz-MS2 transgene, we created a recombinant chromosome containing both. This allowed us to generate a stable fly line with the following genotype: *yw;+;AutoregulatoryEnhancer-Ftz-MS2, Ftz-LlamaTag*. To perform the imaging, we collected virgin females of the genotype *yw; vasa>eGFP; nos>*MCP-mCherry-NLS and *yw; vasa>eGFP/CyO; nos>*MCP-mCherry-NLS. We chose to use a mixture of females with 1 or 2 copies of eGFP as a consistency check in the analysis. This would ensure the background value determined should fall into two groups with the mean being a different by a factor of two which was the case (see below). These were crossed in a cage with the males containing the recombinant chromosome, *yw;+;AutoregulatoryEnhancer-Ftz-MS2, Ftz-LlamaTag*. This ensured that the resulting embryos contained maternally deposited eGFP, MCP-mCherry-NLS, the tagged Ftz locus, and also the AutoregulatoryEnhancer-Ftz-MS2 transgene.

#### Fushi tarazu Protein and ZebraEnhancer-Ftz-MS2

To simultaneously image Ftz protein and transcription from that ZebraEnhancer-Ftz-MS2 transgene, we used the following approach. Using crosses, we generated mother flies that had the following genotype: *yw; vasa>eGFP; nos>*MCP-mCherry-NLS/ *Ftz-LlamaTag* and *yw; vasa>eGFP/CyO; nos>*MCP-mCherry-NLS/ *Ftz-LlamaTag*. We chose to use a mixture of females with 1 or 2 copies of eGFP as a consistency check in the analysis. This would ensure the background value determined should fall into two groups with the mean being a different by a factor of two which was the case (see below). Virgin females of these flies were collected and crossed in a cage with males of the genotype *yw;+;ZebraEnhancer-Ftz-MS2*, and the resulting embryos were imaged. All the progeny had a copy of the *ZebraEnhancer-Ftz-MS2* transgenes and the 50% of embryos containing the *Ftz-LlamaTag* locus could easily be identified and selected on the microscope by looking for Ftz protein. This procedure ensured that the imaged embryos contained maternally deposited eGFP, MCP-mCherry-NLS, the tagged Ftz locus, and also the ZebraEnhancer-Ftz-MS2 transgene.

#### Fushi tarazu Protein and Engrailed PP7

To image Ftz protein and engrailed transcription, we used fly crosses to generate male flies that contained both tagged loci with the following genotype: *en-PP7/CyO; Ftz-LlamaTag/TM3*. These were then crossed in a cage with virgin females that had a mixture of the following genotypes: *nos>tdTomatoPCP-NLS/CyO; vasa>eGFP/TM3* and *nos>tdTomatoPCP-NLS; vasa>eGFP*. The embryos that contained tagged Ftz could be identified by the presence of the protein pattern and those embryos were chosen for imaging. Only the subset of imaged embryos that later displayed engrailed transcription were then used for analysis. This ensured that the embryos that were imaged contained maternally deposited eGFP, PCP-tdTomato-NLS, the tagged Ftz locus, and also the PP7 tagged engrailed locus.

## METHOD DETAILS

### Cloning and Transgenesis

Details of the creation of the various transgenes can be found in the benchling plasmid files. Briefly, the C terminal fusion of Ftz to the LlamaTag utilizing poly-glycine linker was based on a previously published mini-gene^26^. For engrailed, 24 copies of PP7 loop sequences were inserted into the first intron in a poorly conserved region. Both donor constructs for CRISPR-Cas9 mediated homologous recombination were constructed using Gibson Assembly and the gRNAs were selected using the FlyCRISPR website; see sequence repository for further details. Injections were performed by BestGene into strain 54591. A 3xP3 driven dsRed cassette was added to the constructs for visual screening of CRISPR transformants. The autoregulatory and zebra constructs were made in the pBPhi backbone using Gibson Assembly. MS2 loops were placed in the endogenous Ftz intron as done in^26^. The constructs were integrated on chromosome 3 using Bloomington strain 9750.

### Embryo preparation for live imaging

Embryos were dechorionated using bleach, mounted between a semipermeable membrane (Biofolie, In Vitro Systems & Services) and a coverslip (1.5, 18 mm × 18 mm), and embedded in Halocarbon 27 oil (Sigma). Flattening of the embryos makes it possible to image a larger number of nuclei in the same focal plane without significantly impacting early development processes.

### Laser scanning confocal microscopy

Embryos were imaged using a Zeiss LSM 980 confocal microscope. Confocal imaging on the Zeiss was performed using a Plan-Apochromat 40×/1.4NA oil immersion objective. GFP and the RFPs were excited with a laser wavelength of 488 nm (∼35 μW laser power) and 561 nm (∼20 μW laser power), respectively. Fluorescence was detected using the Zeiss QUASAR detection unit.

## QUANTIFICATION AND STATISTICAL ANALYSIS

The sections below on ‘Nuclei segmentation and tracking’ and ‘Ftz level and mRNA measurement’ are based on previous work^26^.

### Nuclei segmentation and tracking

The microscopy experiments resulted in movies with 5 dimensions: *xy* images in two channels (Ftz protein and mRNA/nuclear marker) at multiple *z*-planes and timepoints. The mRNA/nuclear channel functions as a reporter of transcription as well as a nuclear marker. CZI formatted files were imported into MATLAB, after which images were rotated to get the stripe pattern aligned in vertical direction. If the first few timepoints contained partially re-formed nuclei after nuclear division, these timepoints were removed from the movie. To detect and segment the nuclei at each timepoint, a 10 slice sub-stack around the middle of the complete nuclear channel *z*-stack was maximum-intensity projected onto a single plane. Using a sub-stack for the projection was a compromise between maximising detection of all nuclei, even if they moved out of plane, and reduction of motion-blur-induced deformation of nuclei after gastrulation. The MAX-projection was filtered using a Laplace of Gaussians filter with a user-chosen filter size appropriate to enhance features of a size similar to the nuclear diameter. A threshold was applied to the resulting image using Otsu’s method and the segmented nuclei were thickened by one pixel to smooth the edges. At this stage the nuclei were segmented but remained untracked through time. Tracking was done separately in ImageJ/Fiji^47^ using the TrackMate plugin^48^ with the Kalman tracking algorithm (LoG filter = 20 pixel object diameter; Initial search radius = 10 pixel; Search radius = 20 pixel; Max frame gap = 0 frames). The Kalman algorithm is especially suited to objects which move directionally through the frame. The tracks were imported into MATLAB and combined with the previously segmented nuclei, leading to segmented nuclei tracked through time. These segmented areas correspond to the central area of the nuclei, and in order to segment the full nucleus and also to include part of the cytoplasm in which the nucleus resides, the segmented nuclei were dilated until a plane-covering tiling was obtained. From this quasi-hexagonal tiling, the nearest neighbours of each nucleus were determined at every timepoint.

### Ftz level and mRNA measurement

To calculate the Ftz protein level in each nucleus, it is necessary to decide what *z*-slice is used for this measurement. Especially towards to end of nuclear cycle 14, some nuclei start to move in the *z*-direction, meaning that it is not possible to select a single *z*-slice which can be applied for all nuclei. Therefore, we selected per nucleus the brightest slice in the mRNA/nuclear marker channel, which corresponds to the *z*-plane in which the nucleus is in focus at that particular timepoint. The nuclear Ftz level was then calculated as the mean of the protein channel levels within the segmented nucleus area in three slices centred around the selected *z*-plane. mRNA spots were segmented in each *z*-plane by first applying a Laplace of Gaussians filter with a size appropriate to make dot-sized features stand out, followed by binarization with a user-chosen threshold. The choice of a particular value for this threshold was a compromise between maximizing the number of identified ‘real’ dots and minimizing the number falsely identified ‘spurious’ dots. A two-dimensional Gaussian function was fitted to the centre of each detected dot, in order to subtract the local background. From a single mRNA dot, there might be signal in multiple *z*-planes, meaning that the dots belonging to a single mRNA spot must be clustered. From this cluster of dots, the brightest was chosen to represent the intensity of the mRNA spot in question. Finally, this information was linked to the expanded nucleus area in which the spot resides, leading to an mRNA signal per nucleus tracked over time.

### Timepoint interpolation and smoothing

In order to have a consistent time interval between each time point across all embryos, we linearly interpolated the Ftz and mRNA signals to have a 20 s interval. Afterwards we averaged each nuclear Ftz trace with a 4 min moving window centred around each timepoint. Derivatives and second derivatives were also smoothed with a 4 min moving centred window after their calculation. Due to embryo movement, some nuclei moved out of the field-of-view during the movie. Therefore, only those nuclei which were tracked from the start until the gastrulation time were taken into consideration for further analysis.

### Boundary cell classification

To determine the set of AB and PB nuclei in each stripe, we used the stripe pattern at the gastrulation time as a basis. All visible nuclei were first sorted into two categories, ‘stripe’ and ‘non-stripe’ nuclei based on their measured Ftz level at the gastrulation time. The threshold for this sorting was determined by Otsu’s method, leading to the brightest nuclei to be classified as ‘stripe’ nuclei and the dimmest as ‘non-stripe’ nuclei (**Fig S2a**). Using this classification as a basis, the PB and AB nuclei were successively determined. The PB nuclei were defined as those ‘stripe’ nuclei, whose ‘non-stripe’ neighbours are all anteriorly located. Once the PB nuclei were identified, the AB nuclei were defined as those ‘non-stripe’ nuclei, which are anterior neighbours of the PB nuclei (**Fig S2b**). Finally, the PB and AB classified nuclei were assigned to a stripe number. This was done by thresholding a maximum-intensity projection (of an 8-slice sub-stack: 2 slices below to 5 slices above the centre slice) at gastrulation time using Otsu’s method, resulting in a labelled binary image (**Fig S2c-i**). Each nucleus was assigned the label of the nearest region in the labelled image (**Fig S2c-ii**). Finally, the nuclei lists were restricted to those nuclei which are visible from the start until the gastrulation time. The aforementioned steps constituted the first automated guess of the AB and PB nuclei. Since the first guess usually was not entirely accurate, two rounds of refinement were performed. The first round consisted of two diagnostic steps, leading to a list of suggested nuclei which might be misclassified as ‘non-stripe’ nuclei. In the first diagnostic step, for each ‘non-stripe’ nucleus it was determined if its Ftz level increased *after* the gastrulation time, indicating that it might have been misclassified based on its Ftz level *at* gastrulation time. In the second diagnostic step, for each AB nucleus it was checked if it was more than one standard deviation brighter than the average AB nucleus at gastrulation time. The suggested nuclei from both steps (**Fig S2d-i**) were manually checked by inspecting the movie after gastrulation time and the Ftz level traces (**Fig S2d-ii**). If the suggestions were indeed found to be valid, the nuclei were moved from the ‘non-stripe’ list to the ‘stripe’ list, after which the algorithm, as detailed above, was repeated up to the stripe number assignment (**Fig S2e**). For the second round of refinement, only the second diagnostic step was performed, leading to another list of suggested nuclei (**Fig S2f-i/ii**). In addition, the traces of the current AB and PB nuclei were checked for obvious errors, and added to the list of suggestions (**Fig S2f-iii**). This list of suggestions was again manually curated after which one last pass of the algorithm was done up to the stripe number assignment, resulting in the final determination of the AB and PB nuclei in each stripe (**Fig S2g** and **Fig 2a**).

### Switch time and transition time determination

Given that PB nuclei at the end of the movie have a higher Ftz level than the AB nuclei, the divergence time of a PB nucleus was defined as the last timepoint when the Ftz level of that nucleus crossed with the mean Ftz level of its AB neighbours (**Fig 2d**). The transition time was then determined by finding the last timepoint before the divergence time at which the derivative of the PB nucleus Ftz level crossed with the mean of the derivatives of its AB neighbour’s Ftz level (**Fig 2e**). It is possible that the transition time and/or the divergence time occur before the start of the movie, in which case the timepoint was assigned to the first timepoint and was not considered for further analyses involving the transition time. Similarly, it is possibly that the transition and/or divergence time occurs after the gastrulation time. In this case, the nucleus was also not considered for analysis involving transition times, unless a nuclear motion correction was applied (see below).

### Background subtraction and motion correction

#### Autoregulatory and zebra embryos

Autoregulatory enhancer and zebra embryos can display two values of background eGFP levels (high and low), depending on if the mother had one or two copies of eGFP. To determine the values of these levels, three background construct embryos for each background level were imaged and the eGFP levels in all nuclei were analysed (**Fig S1a-i/ii**). Since the eGFP levels over time are not strictly constant, possibly due to some low level of bleaching, the value at the gastrulation time was chosen as the background level. The next step was to assign a particular autoregulatory enhancer or zebra embryo to either the high or low background value. If the median of the Ftz levels in AB nuclei at the gastrulation time was greater than the upper bound (mean + standard deviation) of the high background value, the embryo was assigned the high background value, otherwise the low background value was assigned (**Fig S1b-i/ii, Fig S1c**). To verify that the background subtraction was done correctly, it was checked if the distribution of Ftz levels in the anterior neighbours of AB nuclei at the end of each movie, and after background subtraction, was centred around zero (**Fig S1d**). At this timepoint, these nuclei should contain the lowest amount of Ftz, and are therefore a good proxy for a zero-point.

#### Engrailed embryos

For the engrailed embryos it was not possible to follow the same background subtraction procedure as for the autoregulatory enhancer and zebra embryos (see above). This is due to the blue shifted emission of tdTomato relative to mCherry which meant there was a small amount of bleed though from the red channel to the green. Therefore the mean of the Ftz levels in the anterior neighbours of AB nuclei at the end of each engrailed movie were used as the background value for that particular embryo (**Fig S7a-c)**

Most engrailed transcription happens around or after the gastrulation time (**Fig S6e/g)**. Since many nuclei move in the *z*-direction and out of plane in that time period, the Ftz levels in those nuclei might be altered due to distortions in the point spread function. In order to make statements about the Ftz levels at which *engrailed* transcription is activated, it is therefore necessary to compensate for this nuclear movement. This was done by correcting the Ftz values over time in each nucleus using the values of the mRNA/nuclear marker channel in that nucleus normalized to the initial timepoint. We verified the validity of this procedure by inspecting the eGFP and mRNA/nuclear marker levels in an engrailed background construct. As the eGFP and mRNA/nuclear levels in these constructs should be constant, any deviation from the initial value can be attributed to either bleaching or nuclear movement. By applying the correction as described above, a flat eGFP background level trace was recovered, compensating for the effects of movement after gastrulation and partially for the effects of bleaching before (**Fig S7d/e)**.

### mRNA trace treatment

To reduce the effects of intermittent detection of an mRNA spot (*e*.*g*., the dot moving out of plane momentarily), dips in an mRNA trace of one or two time-frames were ‘filled up’, using a dilation followed by an erosion step. Both the dilation and the erosion used a three time-frame structuring element. This treatment was applied to all constructs. For autoregulatory constructs an additional data treatment step was performed, namely that only persistent mRNA bursts were taken into consideration for analysis. We defined a persistent burst as a burst lasting for at least 3 min based on how long it takes to transcribe the reporter (**Fig S4)**.

### Re-alignment of traces to the transition time

In order to notice common features in traces of PB nuclei related to their developmental trajectory, it is necessary to align the traces according to a fixed point in that trajectory, rather than to an embryo-wide timepoint, such as the gastrulation time. This alignment can be done not only on Ftz traces and their derivatives, but also on mRNA traces, or on functions of these traces, such as the difference in mRNA between neighbours (**Fig 3e/f/g, Fig 4g, Fig 5d/e** and **Fig S4b/c/e/f**). To achieve this re-alignment, the transition time of each PB nucleus was set at t = 0 and the values of that nucleus’ Ftz or mRNA trace were shifted accordingly (**Fig 3d)**. AB neighbours of the PB nuclei were also aligned, by shifting the mean trace of the AB neighbours by the same amount as the PB nucleus in question. Only PB nuclei with a well-defined transition time can be re-aligned. This means that engrailed PB nuclei with a transition time at the first or last time point, and zebra and autoregulatory enhancer nuclei with a transition time at the first timepoint or after the gastrulation time were excluded from this analysis. After all PB and AB neighbour traces of an embryo were re-aligned, the mean of the re-aligned traces was calculated. The embryo-level mean was then used to calculate a ‘construct mean’, namely the mean of all embryo-level means of the construct in question. The number of nuclei participating in this mean varies across the re-aligned time axis, since each nucleus in an embryo has a transition time at a different timepoint. Similarly, the number of embryos participating in the mean at each timepoint might vary, since the movie length of each embryo is different (**Fig S3g/h/i**).

### Ftz level and timepoint shuffling scheme

For each embryo and in each stripe a list of PB nuclei displaying transcription was made. For each nucleus in this list, two numbers were obtained, namely the Ftz level at the start of the first mRNA burst and the timepoint of the start of that burst (relative to gastrulation). This generated, for each embryo and each stripe, a list of ‘observed’ Ftz levels and ‘observed’ timepoints. Next, the ‘observed’ lists of timepoints and the Ftz levels were shuffled, meaning that a random permutation of the elements in the respective list was chosen. Then a list of ‘new’ timepoints was generated from the shuffled list of Ftz levels, and from the shuffled list of timepoints a list of ‘new’ Ftz levels. For the list of shuffled Ftz levels, the procedure was as follows. Each nucleus was assigned a shuffled Ftz value and it was checked at what timepoint that shuffled Ftz level occurred in the Ftz trace of the nucleus (within a 5% tolerance). If there were multiple possible timepoints, a random one was selected from the possibilities. If no possible timepoint was found, then no timepoint was registered for that nucleus. In this manner, list of ‘new’ timepoints was built for each permutation round. The process was repeated until the maximum number of permutations was reached, or 10000 times, whatever number was smaller, in the end resulting in an embryo-level list of ‘new’ timepoints. On the other hand, in the case of the shuffled timepoints, the procedure was as follows. Each nucleus was a assigned a shuffled timepoint and this timepoint was applied to the Ftz trace of the nucleus in question, thereby generating a list of ‘new’ Ftz levels. This process was repeated until the total number of ‘new’ Ftz levels was comparable to the total number of ‘new’ timepoints, in the end resulting in an embryo-level list of ‘new’ Ftz levels. This whole procedure was then applied to all embryos containing a particular stripe, resulting in four stripe-level lists: ‘observed’ Ftz level and timepoint lists, and lists of ‘new’ Ftz levels and timepoints, obtained from the shuffled timepoints and Ftz levels, respectively (Autoregulatory enhancer: **Fig 6f** and **Fig S6b/d**; Engrailed: **Fig 7e** and **Fig S6f/h**). Next, a statistical test was performed to check if the observed and ‘new/shuffled’ lists were the different or not. Since the distributions are non-normal, the non-parametric two-sample Kolmogorov-Smirnov test was chosen (at the 5% significance level), which is both sensitive to location and shape of a distribution. For the null-hypothesis we stated that the ‘observed’ and ‘new/shuffled’ distribution (for either Ftz level or timepoints) were drawn from the same continuous distribution. The alternative hypothesis was that the ‘new/shuffled’ distribution had smaller values (*i*.*e*., earlier timepoints or lower Ftz levels) than the ‘observed’ distribution.

### Notes on histograms and boxplots

For all histograms, the bin width was automatically decided using the Freedman-Diaconis rule, which is less sensitive to outliers in data, and is thus suited for more irregular shaped distributions. In those cases that multiple histograms were plotted in the same figure, the bin width was decided on the histogram with the lowest number of counts. In boxplots, the bottom and top edges of the box denote the 25^th^ and 75^th^ percentiles of the data, respectively. The line inside the box indicates the median, the whiskers extend to the minimum and maximum of the data not considered outliers, and outliers are shown as ‘+’ signs. A data point is considered an outlier if it lies more than 1.5 times the interquartile range from the bottom or top of the box. Notch around the median shows the approximate 95% confidence interval for the median.

### Overview of trace selection and image treatment

General guidelines for trace exclusion are the following. For autoregulatory enhancer embryos, the Ftz and mRNA levels, and associated quantities such as transition time or starting times of the first mRNA burst, are considered not reliable after gastrulation due to nuclear movement, and are therefore disregarded. In case of engrailed embryos, a nuclear movement correction is performed, meaning that for engrailed embryos these values and quantities are included. Additionally, for all embryos, if the transition time cannot be determined, because it is before the start of the movie or after the end, the nucleus in question is excluded if the figure requires analysis of the transition time. Similarly, if an analysis is made of transcription timing and associated Ftz levels, only nuclei are used that actually show transcription within the time period in which we consider the values reliable (see above).

Figure 1c: Maximum intensity projection of green (MaxG) and red channel (MaxR). Gaussian blur applied to MaxR (MaxRBlur). MaxRBlur and MaxG were combined, and relative intensities of channels were adjusted.

Figure 1d: Maximum intensity projection of green (MaxG) and red channel (MaxR). Gaussian blur applied to MaxR (MaxRBlur). MaxRBlur and MaxG were combined, and relative intensities of channels were adjusted.

Figure 1e: Disregard Ftz levels after gastrulation.

Figure 2a/f/g: Maximum intensity projection of green channel. Contrast of channel adjusted.

Figure 2b/c/d/e: Exclude PB nuclei with transition time at/before start of movie or after gastrulation. Exclude PB nuclei without transcription before gastrulation (with mRNA burst length selection). Disregard Ftz levels after gastrulation.

Figure 3a: Exclude PB nuclei with transition time at/before start of movie or after gastrulation. Exclude PB nuclei without transcription before gastrulation (with mRNA burst length selection). Disregard Ftz levels after gastrulation.

Figure 3b/c: Exclude PB nuclei with transition time at/before start of movie or after gastrulation.

Figure 3ef/g; S3: Exclude PB nuclei with transition time at/before start of movie or after gastrulation. Disregard Ftz levels after gastrulation.

Figure 4b: Maximum intensity projection was made of green channel (MaxG). A LoG filter of size comparable to transcription dots was applied to all slices of the red channel, and subsequently a max projection was done (Dots). The Dots image was binarized to retain only the dots (DotsMask). The MaxG and DotsMask image were combined, while adjusting their relative intensities.

Figure 4c/d/e/f; S4: Exclude PB nuclei with transition time at/before start of movie or after gastrulation. Exclude PB nuclei without transcription before gastrulation (with mRNA burst length selection). Disregard Ftz levels after gastrulation.

Figure 4g: Exclude PB nuclei with transition time at/before start of movie or after gastrulation. Apply mRNA burst length selection (with mRNA burst length selection). Only plot the mean at a particular timepoint if more than one embryo is contributing. Disregard mRNA levels after gastrulation.

Figure 5b: Maximum intensity projection was made of green (MaxG) and red channel (MaxR). Gaussian blur was applied to MaxR and the image was normalized to its maximum value (MaxRBlur). Separately, a LoG filter of size comparable to transcription dots was applied to all slices of the red channel, and subsequently a max projection was done (Dots). The Dots image was binarized to retain only the dots (DotsMask). The MaxRBlur and DotsMask images were combined, while adjusting their relative intensities (RedFinal). Lastly, the MaxG was normalized to its maximum value, and subsequently combined with the RedFinal image.

Figure 5c/d; S5a/b/d/e: Exclude PB nuclei with transition time at/before start of movie or after gastrulation. Apply mRNA burst length selection (without mRNA burst length selection). Disregard mRNA levels after gastrulation.

Figure 5e; S5c/f: Exclude PB nuclei with transition time at/before start of movie or after gastrulation. Apply mRNA burst length selection (without mRNA burst length selection). Disregard mRNA levels after gastrulation. Exclude PB nuclei without transcription before gastrulation (with mRNA burst length selection) for autoregulatory enhancer annotation.

Figure 6c/d/e/f, S6a/b/c/d: Exclude PB nuclei with transition time after gastrulation. Exclude PB nuclei without transcription before gastrulation (with burst length selection). Disregard Ftz levels after gastrulation.

Figure 6g-i: PB nuclei that increased their Ftz levels after gastrulation time were manually selected from all available traces. Then it was verified that the nuclear tracking of these nuclei was accurate to exclude an increase of Ftz due to tracking error.

Figure 6g-ii/iii: Maximum intensity projection of green (MaxG) and red channel (MaxR). Gaussian blur applied to MaxR (MaxRBlur). MaxRBlur and MaxG were combined, and relative intensities of channels were adjusted.

Figure 7b: Maximum intensity projection was made of green channel (MaxG). A LoG filter of size comparable to transcription dots was applied to all slices of the red channel, and subsequently a max projection was done (Dots). The MaxG and Dots image were combined, while adjusting their relative intensities.

Figure 7c: Exclude PB nuclei with transition time at/before start of movie or after gastrulation for PB and AB neighbour trace). Exclude PB nuclei without transcription before gastrulation (with mRNA burst length selection) for autoregulatory enhancer annotation. Exclude PB nuclei without transcription (without mRNA burst length selection) for engrailed annotation.

Figure 7d/e; S6e/f/g/h: Exclude PB nuclei with transition time at/after end of movie. Exclude PB nuclei without transcription (without mRNA burst length selection).

Figure S1a/b/c: Disregard Ftz levels after gastrulation time. Only use nuclei with complete trace up to gastrulation.

Figure S2 (images): Maximum intensity projection of green (MaxG) and red channel (MaxR). Gaussian blur applied to MaxR (MaxRBlur). MaxRBlur and MaxG were combined, and relative intensities of channels were adjusted.

Supplemental movie 1: Maximum intensity projection of green (MaxG) and red channel (MaxR). Gaussian blur applied to MaxR (MaxRBlur). MaxRBlur and MaxG were combined, and relative intensities of channels were adjusted.

Supplemental movie 2-5: Only the signal from red channel was used. Signal from the vitelline membrane was removed in each z-slice (if applicable). Maximum intensity projection of red channel followed by Gaussian blur (NucBlur). In parallel, LoG filter (filter size appropriate to enhance dots) was applied to each red channel slice, followed by maximum intensity projection (Dots). For the video the nuclei are shown in Blue and the transcription dots are shown in Yellow. The Dots image was gamma-corrected and intensity scaled to reduce LoG filter artifacts, yielding the Yellow channel. The gamma-corrected and scaled Dots image was subtracted from an intensity scaled NucBlur image, yielding the Blue channel. The combined Yellow and Blue images were automatically rotated to obtain the correct orientation, as needed.

### Supplemental movie titles

[Supplemental movie 1](https://www.dropbox.com/s/vi162b1e6yvixd1/Supplemental%20movie%201.avi?dl=0): Zoom-in movie of Fushi tarazu Protein and MCP-mCherry-NLS (scale bar 20 µm).

[Supplemental movie 2](https://www.dropbox.com/s/74c7jfnxzsgr3au/Supplemental%20movie%202.avi?dl=0): Full embryo movie of Fushi tarazu Protein and AutoregulatoryEnhancer-Ftz-MS2 (mCherry signal only, scale bar 20 µm).

[Supplemental movie 3](https://www.dropbox.com/s/581vsuhbv8p9qsl/Supplemental%20movie%203.avi?dl=0): Zoom-in movie of Fushi tarazu Protein and AutoregulatoryEnhancer-Ftz-MS2 (mCherry signal only, scale bar 20 µm).

[Supplemental movie 4](https://www.dropbox.com/s/1kq3nlkt5efmtfr/Supplemental%20movie%204.avi?dl=0): Full embryo movie of Fushi tarazu Zebra-derived Protein and ZebraEnhancer-Ftz-MS2 (mCherry signal only, scale bar 20 µm).

[Supplemental movie 5](https://www.dropbox.com/s/kc2clzay748u9lx/Supplemental%20movie%205.avi?dl=0): Zoom-in movie of Fushi tarazu Zebra-derived Protein and ZebraEnhancer-Ftz-MS2 (mCherry signal only, scale bar 20 µm).

## Notes

### Competing Interest Statement

The authors have declared no competing interest.

## References

1. Carroll, S. B. Evolution at two levels: On genes and form. PLoS Biol. 3, 1159–1166 (2005).

2. Long, H. K., Prescott, S. L. & Wysocka, J. Ever-Changing Landscapes: Transcriptional Enhancers in Development and Evolution. Cell 167, 1170–1187 (2016).

3. Furlong, E. E. M. & Levine, M. Developmental enhancers and chromosome topology. Science (80-.). 361, 1341–1345 (2018).

4. Gao, T. et al. ScEnhancer: A single-cell enhancer resource with annotation across hundreds of tissue/cell types in three species. Nucleic Acids Res. 50, D371–D379 (2022).

5. Bothma, J. P. et al. Enhancer additivity and non-additivity are determined by enhancer strength in the Drosophila embryo. Elife 4, 1–14 (2015).

6. Weiner, A. J., Scott, M. P. & Kaufman, T. C. A molecular analysis of fushi tarazu, a gene in Drosophila melanogaster that encodes a product affecting embryonic segment number and cell fate. Cell 37, 843–851 (1984).

7. Hiromi, Y. Kuroiwa, a & Gehring, W. J. Control elements of the Drosophila segmentation gene fushi tarazu. Cell 43, 603–613 (1985).

8. Hiromi, Y. & Gehring, W. J. Regulation and function of the Drosophila segmentation gene fushi tarazu. Cell 50, 963–974 (1987).

9. Schier, A. & Gehring, W. Direct homeodomain–DNA interaction in the autoregulation of the fushi tarazu gene. Nature 356, 804–7 (1992).

10. Yu, Y. & Pick, L. Non-periodic cues generate seven ftz stripes in the Drosophila embryo. Mech. Dev. 50, 163–75 (1995).

11. Surkova, S. et al. Characterization of the Drosophila segment determination morphome. Dev. Biol. 313, 844–62 (2008).

12. Frasch, M. & Levine, M. Complementary patterns of even-skipped and fushi tarazu expression involve their differential regulation by a common set of segmentation genes in Drosophila. Genes Dev. 1, 981–995 (1987).

13. Hughes, S. C. & Krause, H. M. Establishment and maintenance of parasegmental compartments. Development 128, 1109–1118 (2001).

14. Lawrence, P. A. & Pick, L. How does the fushi tarazu gene activate engrailed in the Drosophila embryo? Dev. Genet. 23, 28–34 (1998).

15. Lawrence, P. A., Johnston, P., Macdonald, P. & Struhl, G. Borders of parasegments in Drosophila embryos are delimited by the fushi tarazu and even-skipped genes. Nature 328, 440–442 (1988).

16. Dahmann, C., Oates, A. C. & Brand, M. Boundary formation and maintenance in tissue development. Nat. Rev. Genet. 12, 43–55 (2011).

17. Lawrence, P. A. & Struhl, G. Morphogens, Compartments, and Pattern: Lessons from Drosophila? Cell 85, 951–961 (1996).

18. Ingham, P. W., Baker, N. E. & Martinez-Arias, A. Regulation of segment polarity genes in the Drosophila blastoderm by fushi tarazu and even skipped. Nature 331, 73–75 (1988).

19. Lento, W. et al. Wnt / Wingless Signaling in Drosophila Wnt / Wingless Signaling in Drosophila. Cold Spring Harb. Perspect. Biol. 4, 1–16 (2012).

20. Pick, L., Schier, A., Affolter, M., Schmidt-Glenewinkel, T. & Gehring, W. J. Analysis of the ftz upstream element: Germ layer-specific enhancers are independently autoregulated. Genes Dev. 4, 1224–1239 (1990).

21. Schroeder, M. D., Greer, C. & Gaul, U. How to make stripes: deciphering the transition from non-periodic to periodic patterns in Drosophila segmentation. Development 138, 3067–3078 (2011).

22. Clark, E. Dynamic patterning by the Drosophila pair-rule network reconciles long-germ and short-germ segmentation. PLoS Biology vol. 15 (2017).

23. Nasiadka, A & Krause, H. M. Kinetic analysis of segmentation gene interactions in Drosophila embryos. Development 126, 1515–1526 (1999).

24. Han, W., Yu, Y., Altan, N. & Pick, L. Multiple proteins interact with the fushi tarazu proximal enhancer. Mol. Cell. Biol. 13, 5549–5559 (1993).

25. Bothma, J. & Levine, M. Development: lights, camera, action - the Drosophila embryo goes live! ? Curr. Biol. 23, R965–7 (2013).

26. Bothma, J. P., Norstad, M. R., Alamos, S. & Garcia, H. G. LlamaTags: A Versatile Tool to Image Transcription Factor Dynamics in Live Embryos. Cell 173, 1810-1822.e16 (2018).

27. Edgar, B. A., Odell, G. M. & Schubiger, G. Cytoarchitecture and the patterning of fushi tarazu expression in the Drosophila blastoderm. Genes Dev. 1, 1226–37 (1987).

28. Lim, B., Fukaya, T., Heist, T. & Levine, M. Temporal dynamics of pair-rule stripes in living Drosophila embryos. Proc. Natl. Acad. Sci. U. S. A. 115, 8376–8381 (2018).

29. Alon, U. An Introduction to Systems Biology: Design Principles of Biological Circuits. (Chapman and Hall/CRC, 2019).

30. Fukaya, T., Lim, B. & Levine, M. Rapid Rates of Pol II Elongation in the Drosophila Embryo. Curr. Biol. 27, 1387–1391 (2017).

31. Garcia, H. G., Tikhonov, M., Lin, A. & Gregor, T. Quantitative imaging of transcription in living Drosophila embryos links polymerase activity to patterning. Curr. Biol. 23, 2140–5 (2013).

32. Lucas, T. et al. Live imaging of bicoid-dependent transcription in Drosophila embryos. Curr. Biol. 23, 2135–9 (2013).

33. Bertrand, E. et al. Localization of ASH1 mRNA Particles in Living Yeast. Mol. Cell 2, 437–445 (1998).

34. Gratz, S. J., Rubinstein, C. D., Harrison, M. M., Wildonger, J. & O’Connor-Giles, K. M. CRISPR-Cas9 Genome Editing in Drosophila. Curr. Protoc. Mol. Biol. 31.2.1-31.2.20 (2015) doi:10.1002/0471142727.mb3102s111.

35. Heemskerk, J., DiNardo, S., Kostriken, R. & O’Farrell, P. H. Multiple modes of engrailed regulation in the progression towards cell fate determination. Nature 352, 404–10 (1991).

36. Dinardo, S., Sher, E., Heemskerk-jongens, J., Kassis, J. A. & Farrell, P. H. O. Two-tiered regulation of spatially patterned engrailed. 0–5 (1984).

37. Tsai, C. & Gergen, P. Pair-rule expression of the Drosophila fushi tarazu gene: a nuclear receptor response element mediates the opposing regulatory effects of runt and hairy. Development 121, 453–462 (1995).

38. Gregor, T., Tank, D. W., Wieschaus, E. F. & Bialek, W. Probing the limits to positional information. Cell 130, 153–64 (2007).

39. Tkacik, G., Callan, C. G. & Bialek, W. Information flow and optimization in transcriptional regulation. PNAS 105, 12265–70 (2008).

40. Porcher, A. & Dostatni, N. The Bicoid Morphogen System. Curr. Biol. 20, R249–R254 (2010).

41. Fernandes, G. et al. Synthetic reconstruction of the hunchback promoter specifies the role of Bicoid, Zelda and Hunchback in the dynamics of its transcription. Elife 11, 1–32 (2022).

42. Desponds, J., Vergassola, M. & Walczak, A. M. A mechanism for hunchback promoters to readout morphogenetic positional information in less than a minute. Elife 9, 1–56 (2020).

43. Clark, E. & Peel, A. D. Evidence for the temporal regulation of insect segmentation by a conserved sequence of transcription factors. Development (Cambridge) vol. 145 (2018).

44. Soluri, I. V., Zumerling, L. M., Parra, O. A. P., Clark, E. G. & Blythe, S. A. Zygotic pioneer factor activity of odd-paired/zic is necessary for late function of the drosophila segmentation network. Elife 9, 1–36 (2020).

45. Koromila, T. et al. Odd-paired is a pioneer-like factor that coordinates with zelda to control gene expression in embryos. Elife 9, 1–71 (2020).

46. MacArthur, S. et al. Developmental roles of 21 Drosophila transcription factors are determined by quantitative differences in binding to an overlapping set of thousands of genomic regions. Genome Biol. 10, R80 (2009).

47. Schindelin, J. et al. Fiji: An open-source platform for biological-image analysis. Nat. Methods 9, 676–682 (2012).

48. Tinevez, J. Y. et al. TrackMate: An open and extensible platform for single-particle tracking. Methods 115, 80–90 (2017).

49. Fukaya, T., Lim, B. & Levine, M. Enhancer Control of Transcriptional Bursting. Cell 166, 1–11 (2016).

