## Supplemental Information for "Regulating the timing of enhancer transitions is key to defining sharp boundaries of Fushi tarazu expression in the *Drosophila* embryo"

Regulatory landscape of the Ftz locus

Classic studies of the Ftz locus showed that it contains three main enhancers, namely the zebra and upstream enhancers that drive expression in the early embryo, and the neuro enhancer which drives expression in the developing nervous system (**Fig S1**)[1]. These studies also demonstrated that a minigene fragment of the *ftz* locus that includes only these three enhancers are able to make viable flies in a *ftz* null background showing these enhancers are sufficient to enable Ftz to define the body segments and build a functioning nervous system[1,2]. In this study we refer to the upstream enhancer as an autoregulatory enhancer because this enhancer is regulated by the Ftz protein itself[3]. Later studies[4] identified three additional stripe-specific enhancers in the locus connected to each of the the Ftz stripes, except for stripe 4. Since these are not required for rescue, they likely serve as "shadow" enhancers that facilitate patterning robustness in response to stress[5,6]. We think it highly unlikely that other regions exist that regulate Ftz in the early embryo. The region of the genome that regulates Ftz is well defined because the *ftz* gene is flanked on either side by two highly effective and thoroughly characterised insulators[7,8] (SF1 and SF2). Reporter gene studies and sequencing based contact assays have definitively shown that these insulators prevent the *ftz* gene from interacting with sequences outside this insulator-defined region[8–10]. In addition, the chromatin state[11] of the sequences between SF1 and SF2 in the early embryo has been mapped along with the binding of the transcription factor Zelda[12] (which marks active enhancers). Taken together, these results indicate that there are no additional regulatory regions between SF1 and SF2 other than the aforementioned enhancers (**Fig S1**).

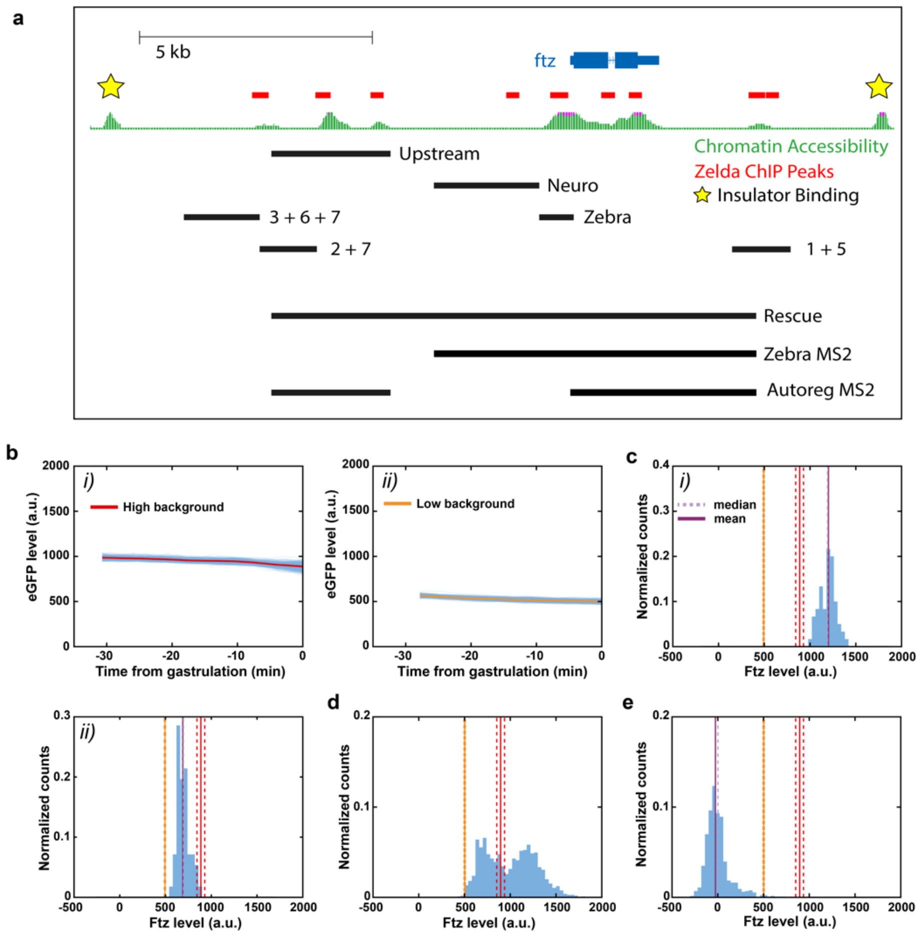


**Figure S1. Regulatory landscape of the *ftz* locus and background subtraction for autoregulatory and zebra enhancer embryos (related to STAR-methods).**

**a.** Schematic illustrating the *ftz* locus along with regions of known regulatory activity, a map of chromatin accessibility, Zelda binding and the location of the insulators. The region capable of rescuing a *ftz*-null mutant as a minigene is also shown together with the sequences used in the autoregulatory and zebra reporters used in this study. **b.** Background eGFP level over time relative to gastrulation for *i)* the high background construct and *ii)* the low background construct. Individual traces are shown including the mean in bold. **c.** Two examples of histograms showing the Ftz level in AB nuclei at the gastrulation time in b *i)* autoregulatory enhancer embryo, and b *ii)* zebra enhancer embryo. **d.** Histogram with Ftz levels in the AB nuclei at the gastrulation time for all embryos of the autoregulatory and zebra enhancer constructs. Note that there are two peaks in the distribution, corresponding to the low and high background values. **e.** Histogram with Ftz levels, after the background correction, in the anteriorly positioned neighbours of the AB nuclei at the final timepoint in each movie, for all embryos of the autoregulatory and zebra enhancer constructs. Note that the background-corrected distribution, and its mean and median, are centred around zero. In panels c-e the low and high background levels are annotated with orange and red lines, respectively. The mean of the background level is a continuous line, whereas a dashed line shows the standard deviation ($N_{embryo} = 3$ for both low and high background). The mean and median of the Ftz level distributions are annotated with dashed and continuous purple lines, respectively.

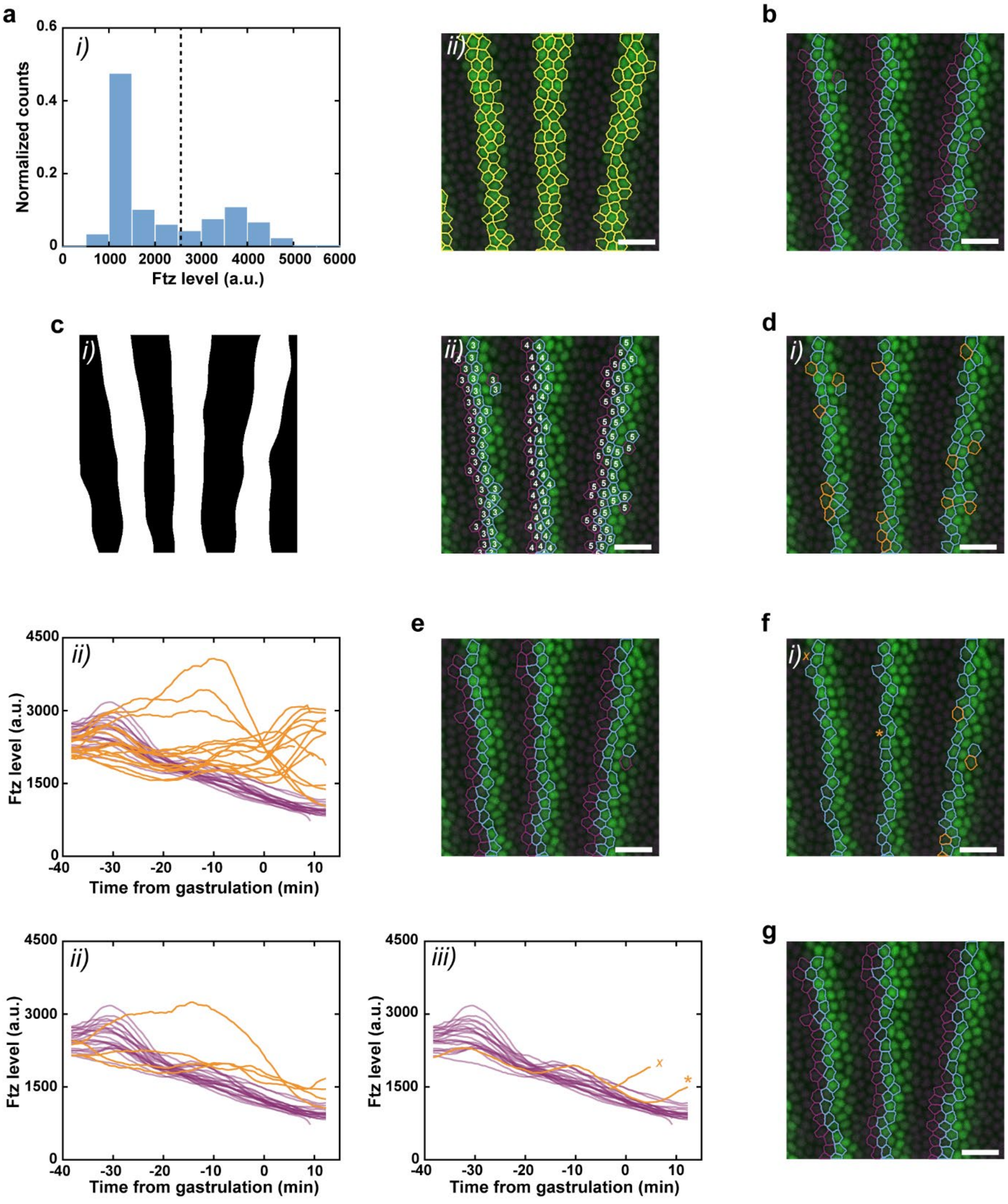


**Figure S2. The algorithm to determine the Anterior and Posterior Boundary (AB and PB) nuclei (related to STAR-methods and Fig 2a)**

**a-***i*) Histogram of Ftz levels of all visible nuclei at gastrulation time, included the threshold on basis of which the nuclei are sorted in 'stripe' and 'non-stripe' nuclei. -*ii*) The initial guess of 'stripe' nuclei. **b.**

The result of the first round of AB and PB nuclei classification. **c-***i*) Thresholded image of the stripe pattern at gastrulation time. -*ii*) Stripe numbering of the initial guess of AB and PB nuclei. **d.** First round of refinement. -*i*) List of suggested nuclei, which might be 'stripe' nuclei instead of 'non-stripe' nuclei. -*ii*) Ftz level traces of these suggested nuclei, together with traces of AB nuclei. **e.** The second guess of AB and PB nuclei. **f.** Second round of refinement. -*i*) List of suggested nuclei, which might be 'stripe' instead of 'non-stripe' nuclei. Also, annotated are two nuclei (x and *) which were missed by the automated suggestions. -*ii*) Ftz level traces of the automatically suggested nuclei, together with traces of AB nuclei. -*iii*) Ftz level traces of the manually suggested nuclei (corresponding to x and * in **f**-*i*), together with traces of AB nuclei. **g.** The final classification of the AB and PB nuclei. For all images: green channel is Ftz and red channel is MCP-mCherry nuclear signal. Scale bars are 20 μm.

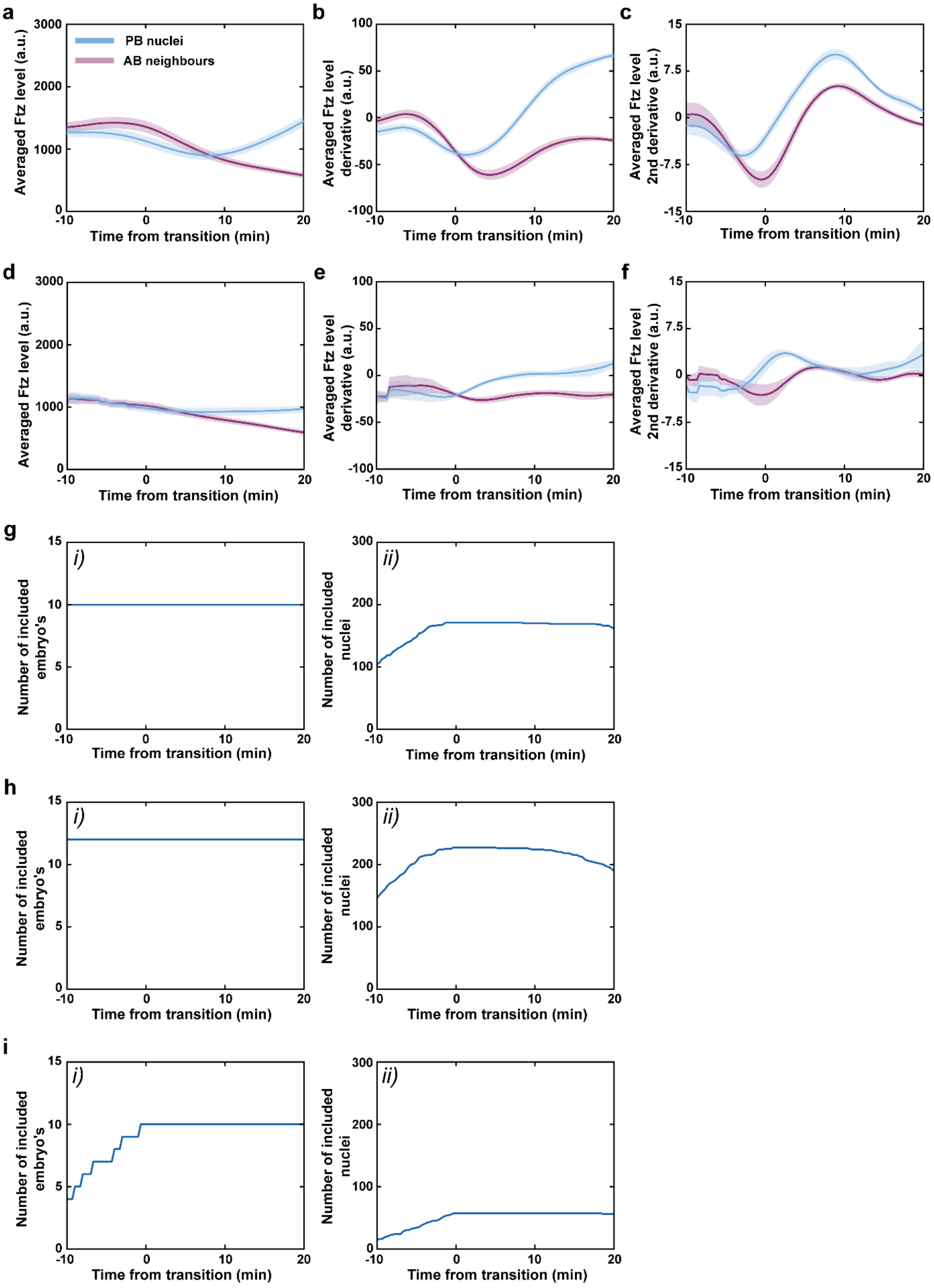


**Figure S3. Averaged Ftz traces (and their derivatives) aligned to the transition time, and the number of embryos and nuclei contributing to these traces (related to STAR-methods and Fig 3).**

Averaged Ftz level aligned to the transition time for PB nuclei and their AB neighbours in stripe 3 (**a**) and stripe 5 (**d**). Note that the Ftz levels cross after the transition time. Averaged Ftz level derivative aligned to the transition time for PB nuclei and their AB neighbours in stripe 3 (**b**) and stripe 5 (**e**). Note that the Ftz level derivatives cross at the transition time, as is expected from the definition of the transition time. Averaged Ftz level second derivative aligned to the transition time for PB nuclei and their AB neighbours in stripe 3 (**c**) and stripe 5 (**f**). Note that the Ftz level second derivatives cross before the transition time. The Ftz level second derivatives are a proxy for mRNA production rate. **g**, **h**, **i**-*i)* Number of embryos contributing to the traces in **a-f** at each timepoint relative to the transition time in stripe 3 (**g**-*i*), stripe 4 (**h**-*i*), stripe 5 (**i**-*i*). **g**, **h**, **i**-*ii)* Number of nuclei contributing to the traces in **a-f** at each timepoint relative to the transition time in stripe 3 (**g**-*ii*), stripe 4 (**h**-*ii*), stripe 5 (**i**-*ii*).

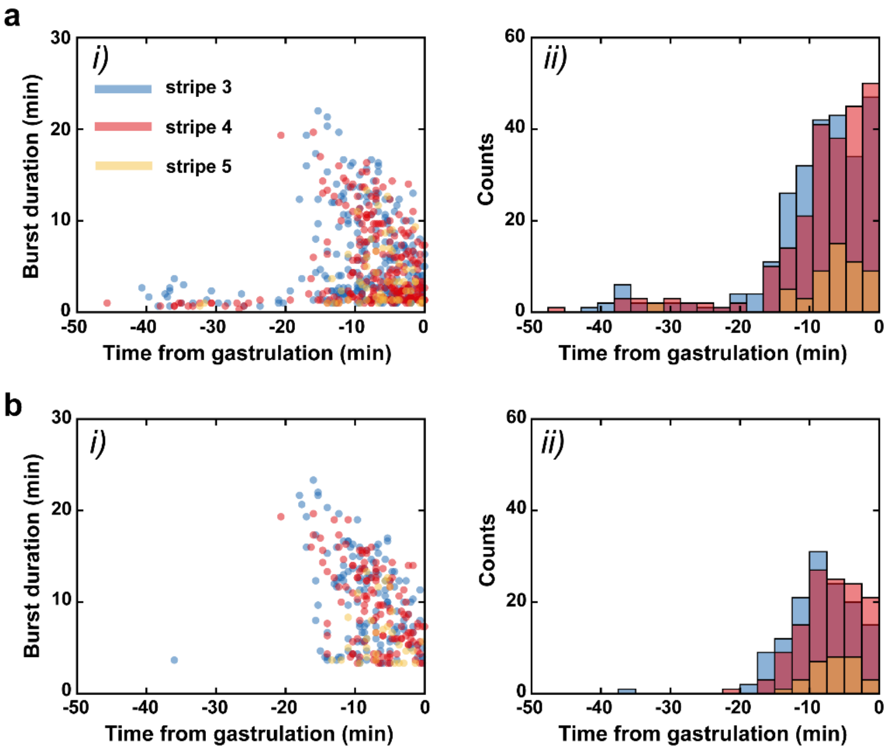


**Figure S4. Persistent mRNA bursts of the autoregulatory enhancer occur only later in nuclear cycle 14 (related to STAR-methods and Fig 4).**

Duration and occurrence of all mRNA bursts in autoregulatory enhancer embryos, displayed per stripe. **a-***i* (before persistent burst selection) and **b-***i* (after persistent burst selection). Scatter plot with all mRNA burst durations over time relative to gastrulation. **a-***ii* (before persistent burst selection) and **b-***ii* (after persistent burst selection). Histogram with number of mRNA bursts over time relative to gastrulation.

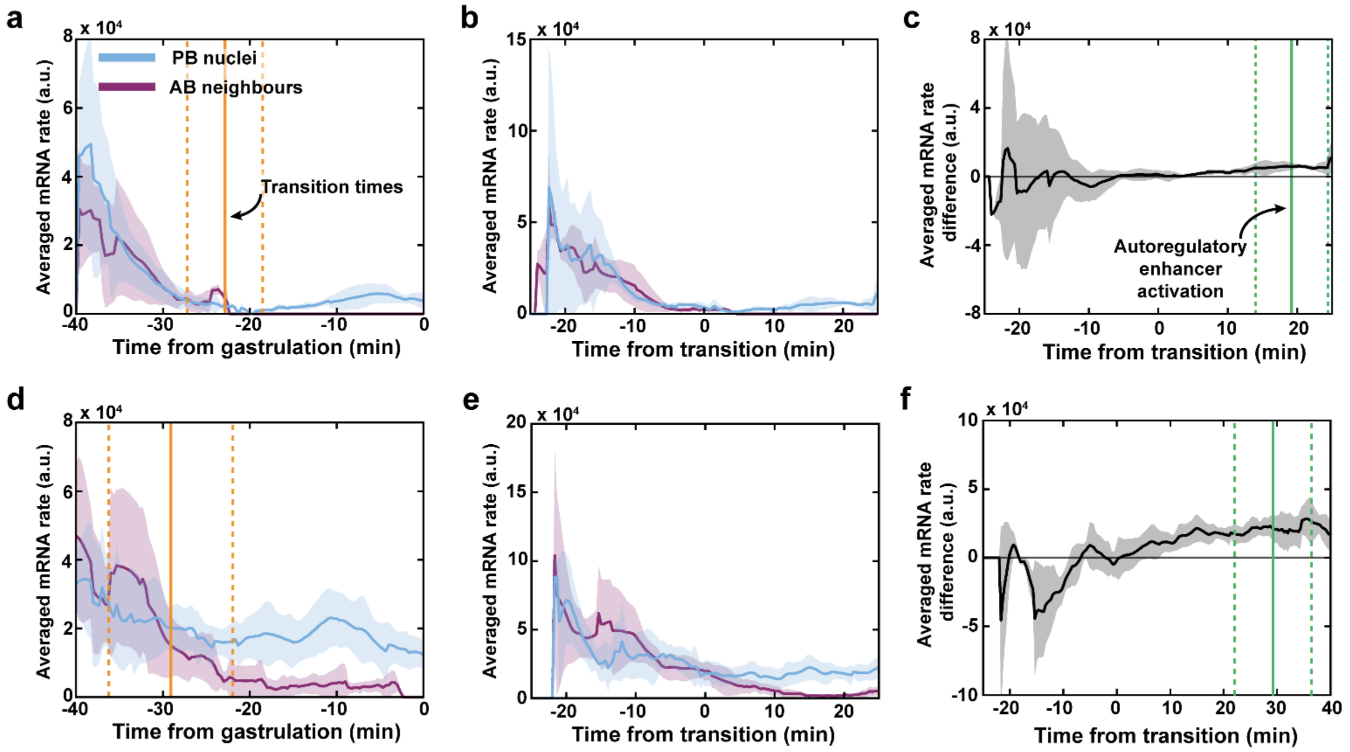


**Figure S5. Comparison of mRNA rate in PB nuclei and their AB neighbours for zebra enhancer embryos (related to Fig 5).**

**a** (stripe 3) and **d** (stripe 5). Averaged mRNA rate in PB nuclei and their AB neighbours in zebra enhancer embryos, relative to gastrulation time. The transition times are annotated using orange lines (continuous for the mean and dashed for the standard deviation). **b** (stripe 3) and **e** (stripe 5). Averaged mRNA rate in PB nuclei and their AB neighbours in zebra enhancer embryos, relative to the transition time. **c** (stripe 3) and **f** (stripe 5). Averaged difference in mRNA rate between PB nuclei and their AB neighbours, relative to transition time. The activation times of the autoregulatory enhancer are annotated using green lines (continuous for the mean, and dashed for the standard deviation).

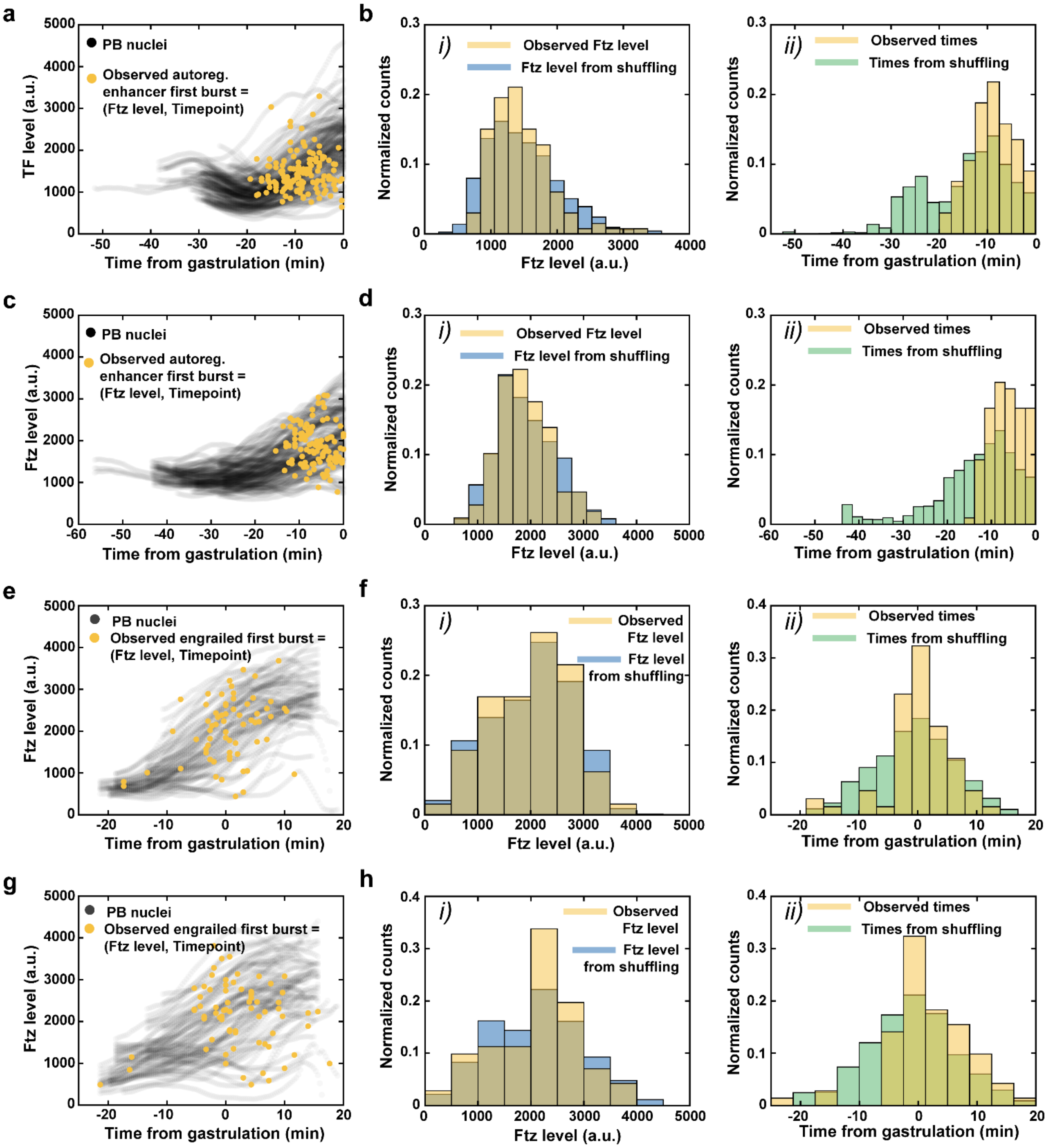


**Figure S6. The Ftz levels and times at which the first mRNA burst in autoregulatory enhancer and engrailed embryos takes place (related to STAR-methods, Fig 6 and Fig 7).**

**a** (stripe 3, autoregulatory enhancer), **c** (stripe 5, autoregulatory enhancer), **e** (stripe 3, engrailed), and **g** (stripe 5, engrailed). Scatter plot of Ftz levels (black) after the transition time, displayed relative to gastrulation, for all PB nuclei showing either autoregulatory enhancer or engrailed transcription. The Ftz level in each PB nucleus at the start of the first mRNA burst is annotated in yellow (autoregulatory enhancer and engrailed). **b** (stripe 3, autoregulatory enhancer), **d** (stripe 5, autoregulatory enhancer), **f** (stripe 3, engrailed) and **h** (stripe 5, engrailed) -*i)* Histogram of the observed Ftz levels (yellow) at the start of the first mRNA burst of the autoregulatory enhancer or engrailed. Shuffling of the starting times of the first mRNA burst in each PB nucleus and application of these shuffled times to the Ftz traces of the respective construct, result in a new Ftz level distribution, shown in blue for both constructs. -*ii)* Histogram of the starting times (yellow) of the first mRNA burst of the autoregulatory enhancer and engrailed. Shuffling of the Ftz levels at these times and a search for the timepoints at which the shuffled Ftz levels occur in each PB nucleus trace of the respective construct, result in a new timepoint distribution, shown in green for both constructs.

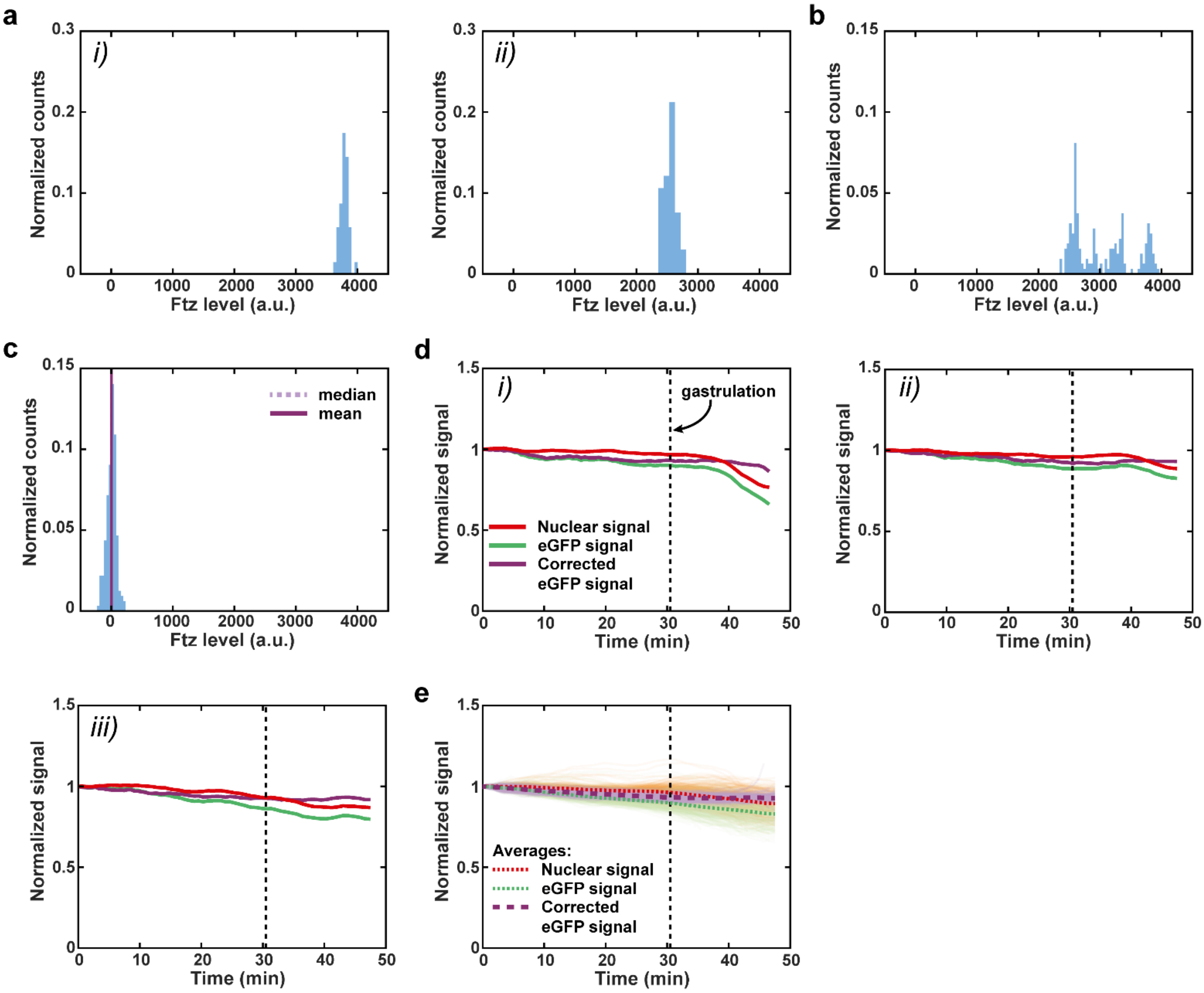


**Figure S7. Background subtraction and nuclear movement correction for engrailed embryos (related to STAR-methods).**

**a.** Histograms of the Ftz levels in the anteriorly positioned neighbours of AB nuclei at the last timepoint of the movie for two engrailed embryos (*i* and *ii*). The mean is used as the background value. **b.** Histogram of the Ftz levels in the anteriorly positioned neighbours of AB nuclei at the final timepoint of each movie, for all engrailed embryos. **c.** Histogram of the Ftz levels, after application of the background correction, in the anteriorly positioned neighbours of AB nuclei at the last timepoint of each movie, for all engrailed embryos. Note that the background-corrected distribution, and its mean and median, are centred around zero. **d** and **e.** Background eGFP (green) and nuclear marker signal (red) for three example nuclei (**d**-*i*, *ii*, and *iii*) and all nuclei (**e**) in an engrailed background embryo, normalized to their respective values at the first timepoint. By using the nuclear marker signal to normalize the background eGFP signal, the purple trace is obtained, which is corrected for nuclear movement and partially for bleaching. The means of the single nucleus traces for eGFP, nuclear marker and corrected eGFP signal are annotated using dashed green, red, and purple lines, respectively. The gastrulation time is annotated with a dashed black line.

**Supplemental Information References**


1. Hiromi, Y., Kuroiwa, a & Gehring, W. J. Control elements of the Drosophila segmentation gene fushi tarazu. *Cell* **43**, 603–613 (1985).

2. Hiromi, Y. & Gehring, W. J. Regulation and function of the Drosophila segmentation gene fushi tarazu. *Cell* **50**, 963–974 (1987).

3. Schier, A. & Gehring, W. Direct homeodomain–DNA interaction in the autoregulation of the fushi tarazu gene. *Nature* **356**, 804–7 (1992).

4. Schroeder, M. D., Greer, C. & Gaul, U. How to make stripes: deciphering the transition from non-periodic to periodic patterns in Drosophila segmentation. *Development* **138**, 3067–3078 (2011).

5. Frankel, N. *et al.* Phenotypic robustness conferred by apparently redundant transcriptional enhancers. *Nature* **466**, 490–3 (2010).

6. Perry, M. W., Boettiger, A. N., Bothma, J. P. & Levine, M. Shadow enhancers foster robustness of Drosophila gastrulation. *Curr. Biol.* **20**, 1562–1567 (2010).

7. Ma, Z. *et al.* Chromatin boundary elements organize genomic architecture and developmental gene regulation in Drosophila Hox clusters. *World J. Biol. Chem.* **7**, 223 (2016).

8. Calhoun, V. C. & Levine, M. Long-range enhancer-promoter interactions in the Scr-Antp interval of the Drosophila Antennapedia complex. *Proc. Natl. Acad. Sci. U. S. A.* **100**, 9878–9883 (2003).

9. Stadler, M. R., Haines, J. E. & Eisen, M. B. Convergence of topological domain boundaries, insulators, and polytene interbands revealed by high-resolution mapping of chromatin contacts in the early Drosophila melanogaster embryo. *Elife* **6**, 1–29 (2017).

10. Batut, P. J. *et al.* Genome organization controls transcriptional dynamics during development. *Science (80-. ).* **375**, 566–570 (2022).

11. MacArthur, S. *et al.* Developmental roles of 21 Drosophila transcription factors are determined by quantitative differences in binding to an overlapping set of thousands of genomic regions. *Genome Biol.* **10**, R80 (2009).

12. Harrison, M. M., Li, X.-Y., Kaplan, T., Botchan, M. R. & Eisen, M. B. Zelda Binding in the Early Drosophila melanogaster Embryo Marks Regions Subsequently Activated at the Maternal-to-Zygotic Transition. *PLoS Genet.* **7**, e1002266 (2011).
